# Defects in plant immunity modulate the rates and patterns of RNA virus evolution

**DOI:** 10.1101/2020.10.13.337402

**Authors:** Rebeca Navarro, Silvia Ambrós, Anamarija Butković, José L. Carrasco, Rubén González, Fernando Martínez, Beilei Wu, Santiago F. Elena

**Affiliations:** Instituto de Biología Integrativa de Sistemas (CSIC - Universitat de València), Paterna, 46182 València, Spain; The Santa Fe Institute, Santa Fe NM87501, USA

**Keywords:** experimental evolution, generalism, plant immunity, plant-virus interactions, response to infection, specialization, virus evolution

## Abstract

It is assumed that host genetic variability for susceptibility to infection necessarily conditions virus evolution. Differences in host susceptibility can either drive the virus to diversify into strains that track different defense alleles (*e.g*., antigenic diversity) or to infect only the most susceptible genotypes. To clarify these processes and their effect on virulence, we have studied how variability in host defense responses determine the evolutionary fate of viruses. To accomplish this, we performed evolution experiments with *Turnip mosaic potyvirus* in *Arabidopsis thaliana* mutants. Mutant plants had disruptions in infection-response signaling pathways or in genes whose products are essential for potyvirus infection. Genotypes were classified into five phenogroups according to their response to infection. In order to disentangle how host susceptibility affects virus adaptation, independent viral lineages were evolved in selected plant genotypes. Evaluating disease-related traits of the evolved lineages, we found that evolution proceeded faster in the most resistant hosts than in the most permissive ones, as expected for adaptation to a harsh environment. By sequencing the genomes of the evolved viral lineages, we found that the multifunctional protein VPg turned out to be the target of selection in most host genotypes. When all evolved viral lineages were tested for fitness in all plant genotypes used in the experiments, we found compelling evidences that generalist viruses were selected by the most restrictive plant genotypes, while permissive genotypes selected for specialist viruses. Overall, this work illustrates how different host defense signaling pathways constrain not only disease-related traits but virus evolution.

## Introduction

The spectrum of disease severity can be attributed to heterogeneity in virus virulence or in host factors; the two are not necessarily independent explanations and they must actually complement and/or interact with each other resulting in an arms-race coevolutionary process. A problem faced by viruses is that host populations consist of individuals that had different degrees of susceptibility to infection (Schmid-Hempel and Koella 1994; Pfenning 2001; Sallinen et al. 2020). Therefore, adaptive changes improving viral fitness in one host genotype may be selected against, or be neutral, in an alternative one. Genetic variability in susceptibility of hosts and infectiousness of viruses have been well studied in animals and plants (*e.g*., Schmid-Hempel and Koella 1994; Altizer 2006; Hughes and Boomsma 2006; Brown and Tellier 2011; Anttila et al. 2015; Parrat et al. 2016; González et al. 2019; Sallinen et al. 2020). The interaction between host and parasite genotypes has been explained in the light of two different theoretical models that represent the two extremes on a continuum of possibilities. At the one extreme, the so-called gene-for-gene (GFG) model, in which a virus genotype exists that can infect all host genotypes and a universally susceptible host genotype should also exist (Flor 1956). Resistance occurs when a host “resistance” gene is matched by at least one virus’ “avirulence” gene. Polymorphism in infectivity and resistance can be maintained only if virulence pays a cost. At the other extreme, the matching-alleles (MA) model is based on self-versus non-self-recognition systems in invertebrates. Infection is not possible unless the virus possesses all alleles that match those of the host (Frank 1993). In this case, polymorphism in infectivity and resistance are maintained by negative frequency-dependent selection.

Variability within host populations arises from nonrandom spatial distributions of genotypes: social groups of animals are more closely related to each other than to other members of the population, and plant crops are generally cultivated as genetically homogeneous plots. These situations facilitate virus transmission among genetically similar host genotypes. Spatial structure and local migration predict evolution of less aggressive exploitation in horizontally transmitted parasites (reviewed in Parrat et al. 2016). For example, Boots and Mealor (2007) observed that *Plodia interpunctella* granulosis virus evolved in spatially structured lepidopteran host populations became less virulent than the one maintained in well-mixed host populations. Berngruber et al. (2015) showed that a latent *λ* bacteriophage won competitions against a virulent one in a spatially structured host populations but lost in well-mixed populations. Similar results were observed in laboratory evolution experiments with *Arabidopsis thaliana* and tobacco etch virus (TEV): a negative association between the permissiveness to infection of different plant accessions and TEV virulence was observed (Hillung et al. 2014). Finally, host population structure also promotes coexistence of hosts and parasites reducing the pathogens dispersal (Brockhurst et al. 2006).

Theoretically, in the absence of host heterogeneity, parasites must evolve toward a host exploitation strategy that maximizes transmission with low virulence (Haraguchi and Sasaki 2000; Regoes et al. 2000; Rodríguez and Torres-Sorando 2001; Ganusov et al. 2002; Gandon 2004; Lively 2010; Moreno-Gámez et al. 2013). Chabas et al. (2018) experimentally showed that evolutionary emergence is more likely when the host population contains intermediate levels of resistant hosts, confirming this prediction using a set of phages and bacterial hosts with alterations in the CRISPR/Cas-mediated immunity.

Viral infection of plants is a complex system in which the virus parasitizes the host and utilizes all its cellular resources to replicate and systemically spread. In response, plants have evolved intricated signaling mechanisms that limit the spread of the virus, resulting in resistance (Zhou and Zhang 2020). A variety of factors contribute to plant resistance to viral infections. Broadly speaking, these factors can be classified into *basal* if they are pre-existing and limit within-cell propagation and cell-to-cell spread, and *inducible* if they are only activated upon infection and inhibit systemic virus movement and replication. Basal mechanisms include susceptibility (*S*) genes that involve alleles of cellular proteins that do not interact properly with viral factors; *e.g*., translation initiation factors required by the virus for successful exploitation of the cell’s protein synthesis machinery, heat shock proteins that assist the formation of multiprotein complexes, or DNA binding phosphatases (Carr et al. 2010; Mäkinen 2019). In contrast, inducible mechanisms include genes whose expression results in a broad-scale change in plant physiology via diverse signal transduction pathways, particularly those regulated by the hormones salicylic acid (SA), jasmonic acid (JA) and ethylene (ET) (Soosaar et al. 2005; Carr et al. 2010). These changes include local cell apoptosis (*e.g*., hypersensitive responses - HR; Loebenstein 2009), the upregulation of nonspecific responses against many different types of pathogens throughout the entire plant (systemic acquired resistance - SAR- and induced systemic resistance - ISR) (Kachroo et al. 2006; Carr et al. 2010), and the activation of the RNA-silencing-based resistance, which seems to play a role both in basal and inducible mechanisms (Voinnet 2001; Carr et al. 2010). Early host responses following virus detection include changes in ion fluxes (mainly Ca^++^), activation of signaling pathways, major alterations of transcriptomic profiles, generation of reactive oxygen species (ROS) and production of nitric oxide (NO) (Soosaar et al. 2005). These immediate changes are followed by HR and the recruitment of SA and JA/ET signaling pathways. The SA-mediated defense signaling pathway results in SAR, while the JA/ET-mediated defense signaling pathway results in ISR, the latter being specifically involved in interactions between plants and beneficial microbes. Indeed, it appears that ISR is not effective against infection by most viruses (Ton et al. 2002; Loebenstein 2009; Pieterse et al. 2009). Both SAR and ISR pathways converge into two master regulators, the *ENHANCED DISEASE SUSCEPTIBILITY 1* (*EDS1*) and the *PHYTOALEXIN DEFICIENT 4* (*PAD4*) genes; *EDS1* and *PAD4* repress ISR and promote SAR. Although the SA and JA/ET pathways have been classically viewed as mutually antagonistic, several studies have revealed positive and negative crosstalk between them (van Wees et al. 2000; Pieterse et al. 2012) as well as with the RNA-silencing pathway (Soosaar et al. 2005; Carr et al. 2010; Yang et al. 2020). This crosstalk, which has the network topological structure of an incoherent feed-forward loop, creates robustness and tunability in the plant immune network (Mine et al. 2017).

An open question is how the intricacies of the underlying defense regulatory pathways of *A. thaliana*, the existence of multiple defense responses and their potential crosstalk determines the evolutionary fate of viruses. For instance, would two independent defense pathways select for specialist viruses adapted to counteract each one thus conforming a fitness tradeoff? Would a tradeoff in the investment of plant resources between pathways (*e.g*., SAR *vs* ISR) result in a diversification of virus adaptive strategies? Would viruses targeting crosstalk points across pathways evolve as generalists? Here we will use turnip mosaic virus (TuMV; species *Turnip mosaic potyvirus*, genus *Potyvirus*, family *Potyviridae*) - *A. thaliana* experimental pathosystem to explore these questions. After exploring the variability in phenotypic responses to TuMV infection of a collection of *A. thaliana* mutants in basal and inducible resistance mechanisms, we choose nine mutants which cover the entire spectrum in phenotypic responses. Then, we performed evolution experiments on each plant mutant genotype and track the evolution of several disease-related traits. At the end of the evolution experiment, we evaluated the effect of host genotypes in the rates of virus evolution and the contribution of historical contingency, selection and stochasticity into the outcome of evolution. Next, we explored whether the different TuMV lineages evolved as specialist or generalists depending on the disrupted defense mechanism of their local hosts. Finally, we sought to identify the molecular changes experienced in the genome of the different viral lineages and to explore the possible adaptive value of a few convergent mutations.

## Results

### Classification of *A. thaliana* genotypes according to their phenotypic response to TuMV infection

The 22 *A. thaliana* mutant genotypes used in this study are shown in table 1, including information about the affected signaling pathways or cellular processes as well as the expected phenotype of infection relative to wildtype (WT) plants based on the description of these mutants. Genotypes were classified according to their phenotypic response to TuMV infection based in five different disease-related traits measured 18 days post-inoculation (dpi). The disease-related traits measured were change in dry weight (*ΔDW*), disease progression (area under the disease progress step curve or *AUDPS*), infectivity (*I*), symptoms severity (*SS*), and viral load (*VL*). Figure 1 shows the nearest-neighbor clustering of genotypes according to their multi-trait phenotypic response to TuMV infection (supplementary figure S1, Supplementary Material online); we found five significant phenogroups (hereafter named as G1 to G5). For all four members of G1 the response was consistent with an enhanced resistance response to TuMV infection: no significant *ΔDW* whilst a significant and consistent reduction in *AUDPS*, *I*, *SS*, and *VL* relative to infected WT plants. Three members of G1 had enhanced SAR response (table 1) and the fourth one (*i4g2*) is a well-known *S* gene involved in plant resistance to potyviruses (Nicaise et al. 2007; Gallois et al. 2010). The only member of G2 was the strong apoptosis-inducer *p58^IPK^*, which shows no significant changes in *AUDPS*, *I*, *SS*, and *VL* relative to infected WT plants but a significant reduction in *ΔDW*. Infected *p58^IPK^* plants were heavier than the noninfected controls which points to a case of increased tolerance to infection. The only member of G3 was *dbp2*, that specifically increased resistance against another potyvirus (Castelló et al. 2011). This mutant showed a very interesting response to TuMV infection: *ΔDW* was significantly increased while *AUDPS*, *I* and *SS* were significantly reduced; no effect in *VL* was observed. Thus, *dbp2* represents an intermediate response to TuMV infection between the highly resistant G1 and those genotypes in G4. G4 represents a sort of hotchpotch formed by genotypes that do not show a clear phenotypic difference from WT. Remarkably, though, with the exception of *cpr5-2*, *dcl2 dcl4*, and *dip2*, all show significant reductions in *VL*. Finally, *jin1* was the only member of G5, which shows a significant increase in all measured traits except *VL*, consistent with and enhanced susceptibility to TuMV infection. This is a surprising finding since *jin1* has been described as inducing a constitutive expression of the SA-dependent defenses that made plants extremely resistant to the infection of biotrophic bacterial pathogens (Laurie-Berry et al. 2006). To further characterize the progression of infection, we selected a subset of representatives from groups G1 (*eds8-1*) and G4 (*cpr5-2*, *dcl2 dcl4*, *hsp90-1*, and WT) and evaluated daily *I*, *SS* and *VL* (supplementary figure S2, Supplementary Material online). The *cpr5-2* plants started showing symptoms faster, though only 80% of plants showed symptoms after 15 dpi. Only 20% of *eds8-1* plants showed symptoms of infection, which were delayed. Infected *hsp90-1* and *dcl2 dcl4* plants were undistinguishable from the infected WT ones. Regarding *VL*, all genotypes accumulated less virus than WT plants, amongst them *cpr5-2* accumulated the least amount of virus (supplementary figure S2, Supplementary Material online).

**FIG. 1.**
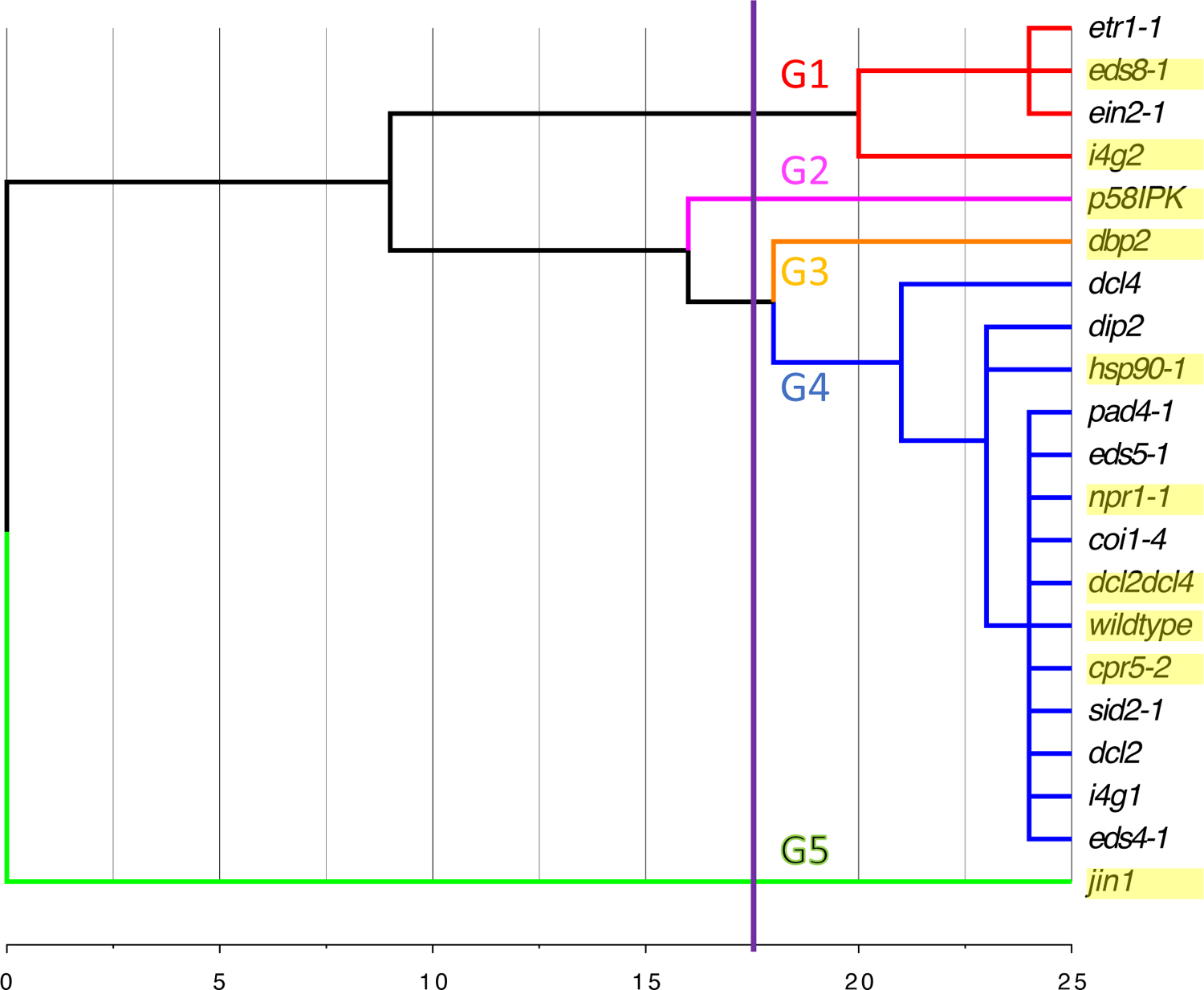
Nearest-neighbor clustering, based on the squared Euclidean distance, of the 21 mutant genotypes of *A. thaliana* according to their phenotypic similarity in response to TuMV infection. Genotypes selected as hosts for the evolution experiments are highlighted in yellow.

**Table 1.**
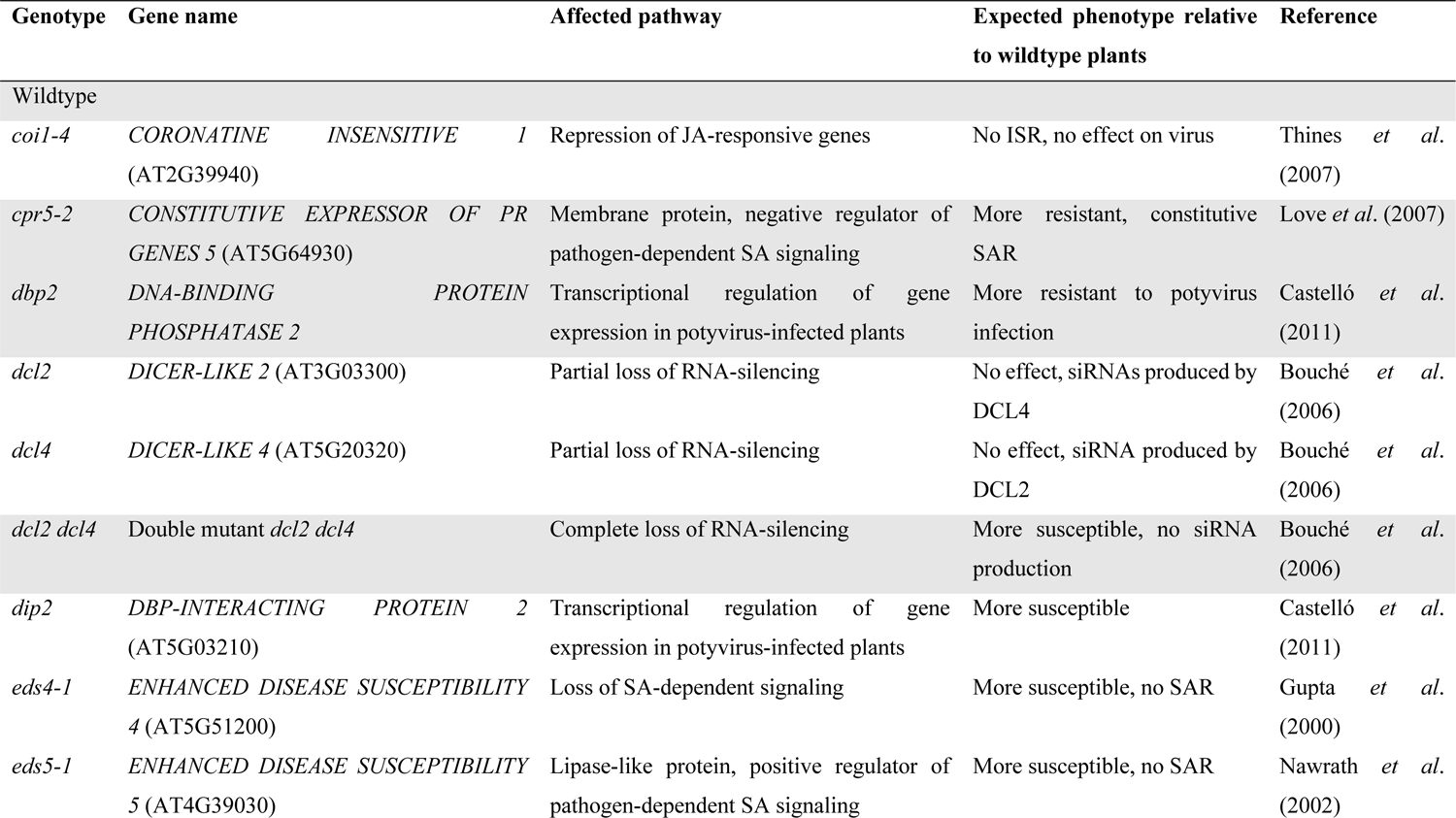
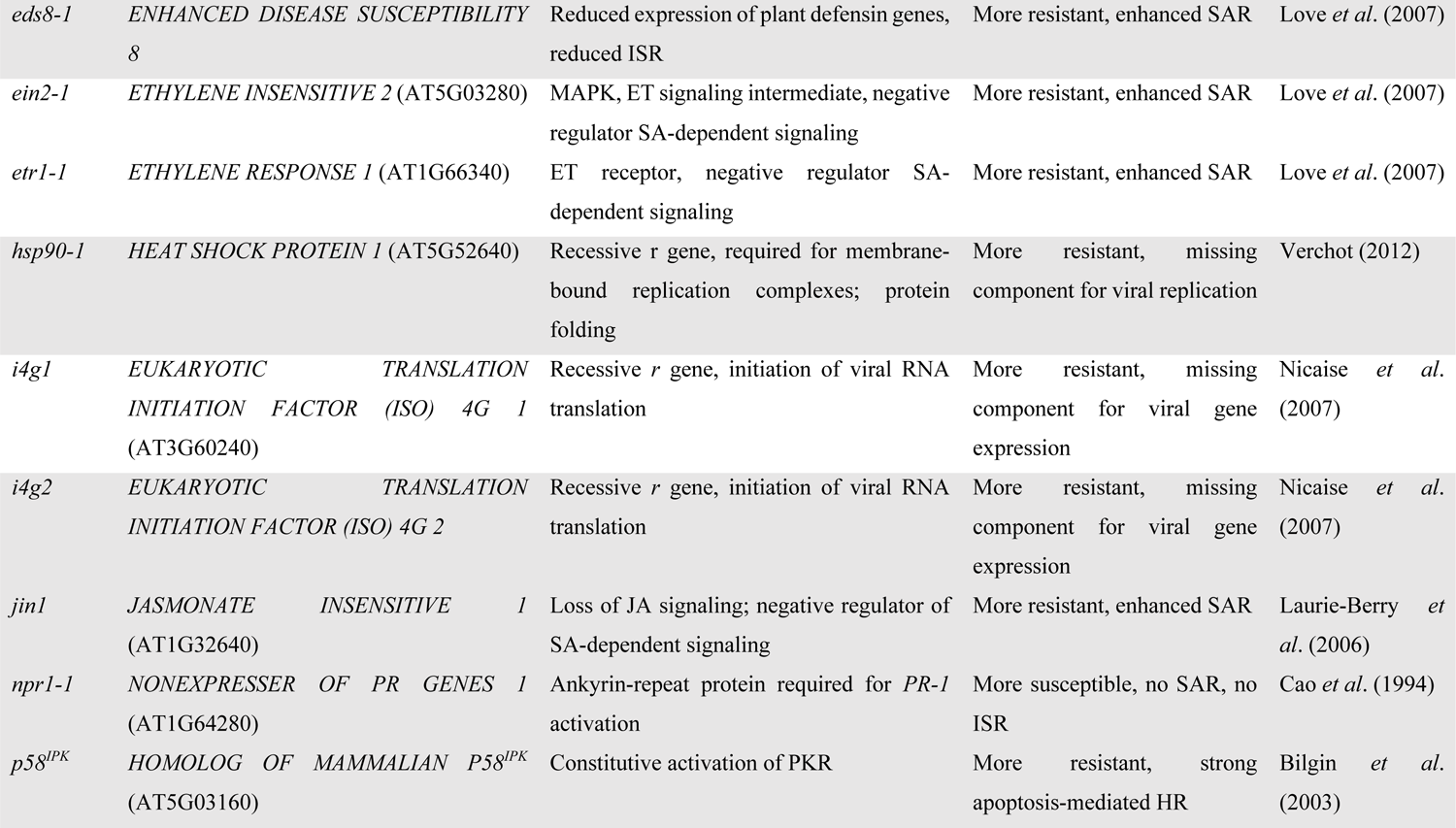
Different arabidopsis genotypes used in this study. Highlighted in gray are those used for the evolution experiments.

Representatives from the five groups were selected for the subsequent evolution experiment (shadowed rows in table 1): *eds8-1* and *i4g2* from G1 representing the hardest selection regimes (more resistant genotypes); *p58^IPK^* from G2; *dbp2* from G3; *cpr5-2*, *dcl2 dcl4*, *hsp90-1*, *npr1-1*, and WT from G4; and *jin1* from G5 representing the softest selective regime (less resistant genotype).

### Experimental evolution in representative *A. thaliana* genotypes

First, we sought to evaluate the dynamics of evolution for *AUDPS* (figure 2), *I* (figure 3) and *VL* (figure 4). Data in these figures were fitted all together into the multivariate analysis of covariance (MANCOVA) model described in equation 1 of the Materials and Methods section. Notice that lineage *cpr5-2*/L2 showed a quite different evolution pattern from the other four lineages evolved in *cpr5-2*. It never increased in either of the three traits and was lost after passage five (figures 2 and 3). Therefore, we ended up with 46 evolved lineages (two in the WT, four in *cpr5-2* and five in each of the other eight plant genotypes).

**FIG. 2.**
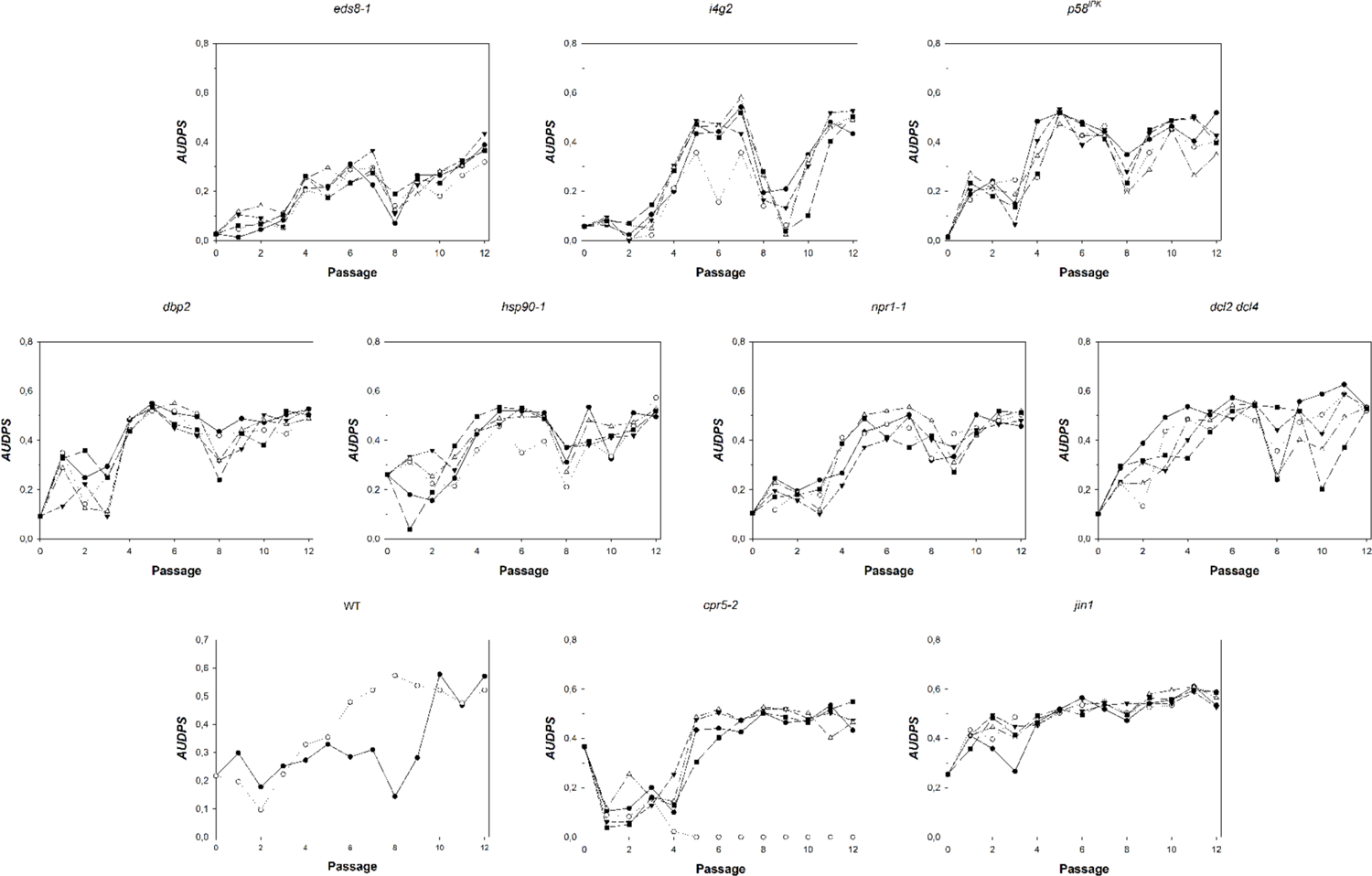
Evolution of disease progression (*AUDPS*) along the serial passages of experimental evolution on each different host genotype. Different symbols and lines represent the independent evolutionary lineages. Panels are arranged from the most resistant genotype (*eds8-1*) to the most sensitive one (*jin1*) according to the groups defined in figure 1.

**FIG. 3.**
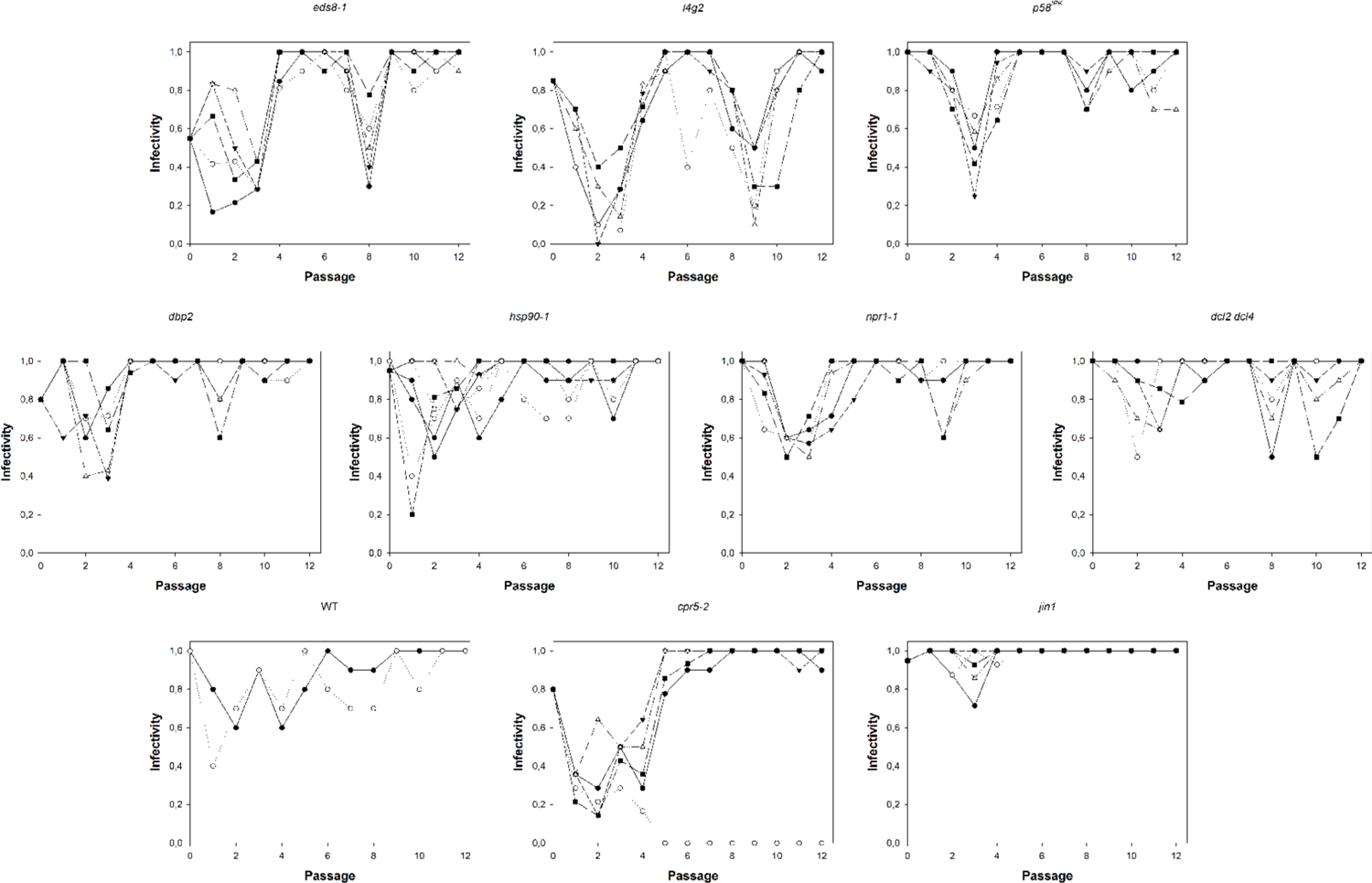
Evolution of infectivity (*I*) along the serial passages of experimental evolution on each different host genotype. Different symbols and lines represent the independent evolutionary lineages. Panels are arranged from the most resistant genotype (*eds8-1*) to the most sensitive one (*jin1*) according to the groups defined in figure 1.

**FIG. 4.**
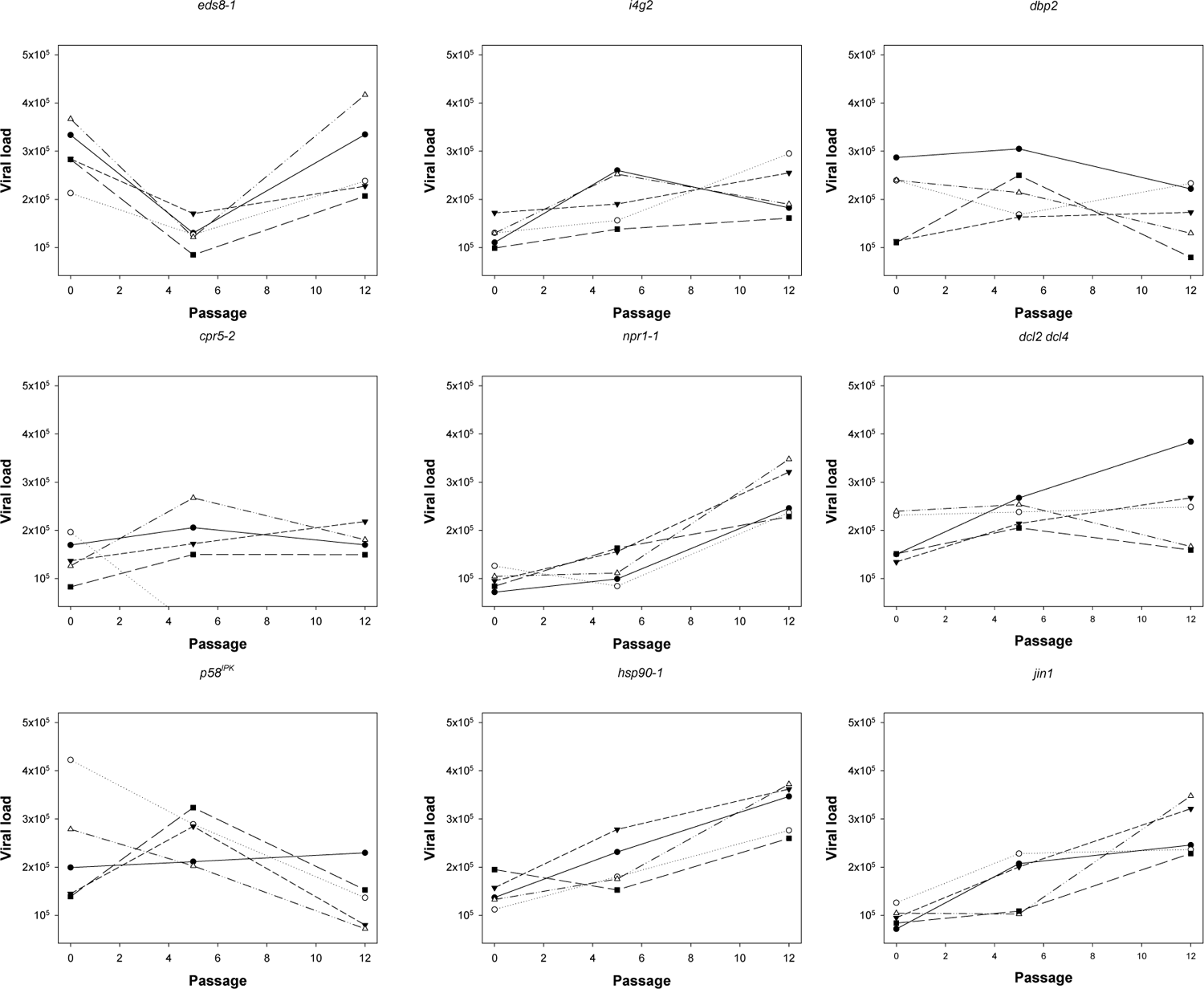
Evolution of viral load (*VL*) along the serial passages of experimental evolution on each different host genotype. Different symbols and lines represent the independent evolutionary lineages. Panels are arranged from the most resistant genotype (*eds8-1*) to the most sensitive one (*jin1*) according to the groups defined in figure 1. TuMV *VL* was not evaluated for WT plants along the evolution experiment.

Table 2 shows the results of this MANCOVA. Despite a considerable amount of noise in the time series, we found some significant results that can be summarized as follows. First, a net significant and large (η^2^_P_ = 0.735) effect associated to the evolutionary passages (*t*) has been observed, indicating that the three phenotypic traits related to TuMV infection evolved during the experiment. Second, plant genotypes (*G*) had a highly significant and large (η^2^_P_ = 0.547) effect on the phenotypic traits, suggesting that TuMV evolutionary dynamics were strongly influenced by the local host genotype in which it was being passaged. More interestingly, this effect was strongly dependent on the passage number (*t×G*; η^2^_P_ = 0.463), thus suggesting that the slope of the relationship between phenotypic traits and time was influenced by the local host genotype. This effect will be further evaluated in the next section. Third, independent lineages evolved in the same host genotype show a strong degree of evolutionary parallelism, as indicated by nonsignificant differences among viral lineages within local host genotype (factors *L*(*G*) and *t×L*(*G*)). However, this last conclusion should be taken with certain caution since the power of the two tests is relatively low (1 *− β* < 0.800) and hence we might be wrongly failing to reject the null hypothesis (type II error).

**Table 2.** Results of the multivariate analysis of variance (MANCOVA) for the three phenotypic traits evaluated along the course of experimental evolution. The different factors are defined in equation 1.

| Source of variation | Wilk's $\Lambda$ | $F$ | $df_1, df_2$ | $P$ | $\eta_p^2$ | $1 - \beta$ |
| --- | --- | --- | --- | --- | --- | --- |
| $\Pi$ | 0.004 | 3377.616 | 3, 42 | $< 0.001$ | 0.996 | 1.000 |
| $t$ | 0.265 | 38.799 | 3, 42 | $< 0.001$ | 0.735 | 1.000 |
| $G$ | 0.093 | 6.467 | 24, 122.414 | $< 0.001$ | 0.547 | 1.000 |
| $t \times G$ | 0.155 | 4.608 | 24, 122.414 | $< 0.001$ | 0.463 | 1.000 |
| $L(G)$ | 0.529 | 0.278 | 108, 126.692 | 1.000 | 0.191 | 0.373 |
| $t \times L(G)$ | 0.466 | 0.351 | 105, 126.677 | 1.000 | 0.225 | 0.487 |

$\eta_p^2$ : magnitude of the effect; $1 - \beta$ : power of the test.

A relevant question in evolutionary biology is the extent in which ancestral differences determine the fate of evolution. In other words, the relative contribution of adaptation, chance and historical contingency to the evolution or organismal fitness. Following the logic exposed by Travisano et al. (1995), we can imagine two situations. First, ancestral statistical differences among phenotypes could be preserved along evolution despite a net increase in the mean trait values (due to selection) and differences among replicated lineages (due to chance; *i.e*., mutation and drift). In this situation, we should expect a nonzero slope in a regression of the evolved phenotypic values against the ancestral ones. The closer the slope to a value of one, the more importance of ancestral differences. Second, if initial trait variation among ancestral genotypes was eliminated from the evolved populations because the combined effect of adaptation and chance, we should expect a regression slope smaller than one. The less effect of ancestral differences, the flatter the slope, being zero in the extreme case where ancestral differences are completely erased. To evaluate the role of the ancestral differences in *AUDPS*, *I* and *VL* of TuMV across the nine different plant genotypes, we evaluated the magnitude of their change at the end of the evolution experiment. Figure 5 shows the plots of evolved *vs* ancestral values. For illustrative purposes, the solid diagonal lines represent the null hypothesis of absolute preservation of ancestral differences. In the case of *AUDPS* (figure 5A), a significant regression exists between evolved and ancestral values (*R* = 0.491, *F*_1,44_ = 14.000, *P* < 0.001), with slope 0.284 ±0.076 (±1 SD). Since the slope is still significantly different from zero, yet clearly flatter than the diagonal (*t*_44_ = 9.422, *P* <0.001), we conclude that ancestral differences have been mostly removed by the combined action of selection and chance, yet not completely erased. In the case of *I* (figure 5B), the regression was not significant (*R* = 0.094, *F*_1,45_ = 0.405, *P* = 0.528), thus suggesting that for this trait ancestral differences among *A. thaliana* genotypes have been fully erased by adaptation and chance. Finally, in the case of *VL* (figure 5C; WT not included in this analysis), the regression of evolved values on the ancestral ones was significant (*R* = 0.417, *F*_1,43_ = 8.822, *P* = 0.005), suggesting again that ancestral phenotypic differences have been largely removed by selection and chance yet some still persist (regression slope *−*0.142 ±0.048 significantly different from one: *t*_42_ = 23.903, *P*< 0.001).

**FIG. 5.**
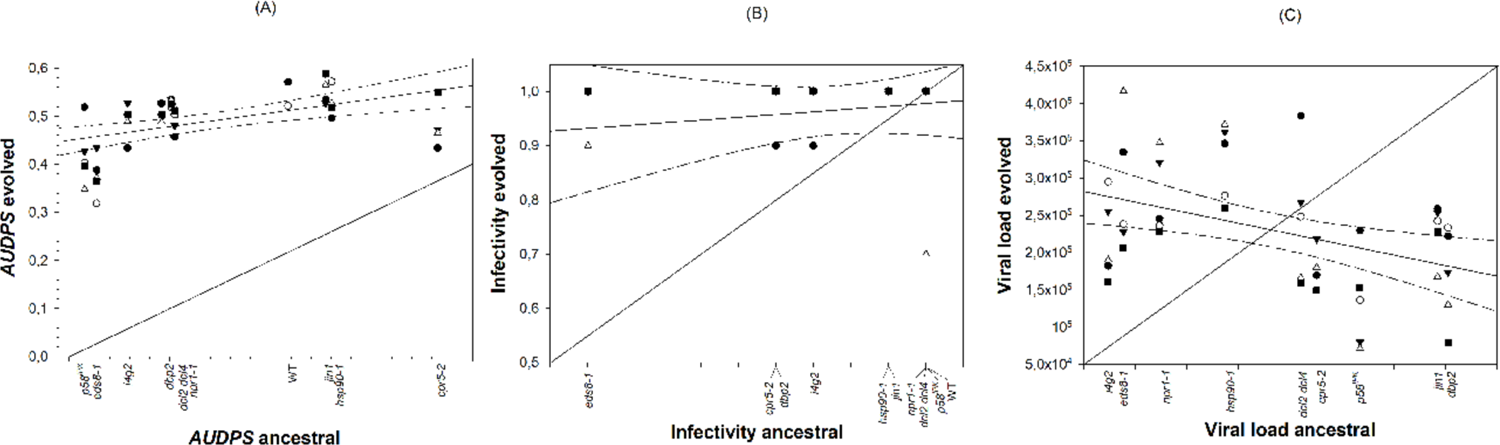
Test of the contribution of historical contingency to the observed pattern of adaptation of TuMV to the different *A. thaliana* genotypes. (A) Evolved *vs* ancestral values for *AUDPS*, (B) for *I*, and (C) for *VL*. The diagonal lines represent the null hypothesis of historical differences being fully preserved despite adaptation. The solid lines represent the linear regression of the data (dashed lines represent the 95% CI for the regression lines).

### Rates of phenotypic evolution for disease-related traits

As mentioned in the previous section, we have observed a significant effect of *A. thaliana* genotypes in the temporal evolution of the three phenotypic traits. To further explore this effect, we estimated the rates of phenotypic evolution of *AUDPS* and *I* using the ARIMA(1,0,0) model described by equation 2 in Materials and Methods. *VL* was not included in this analysis since the number of data points (three) in the time series was the same as the number of parameters to be estimated, thus likely incurring in overfitting problems.

Figure 6 shows the estimated rates of evolution (*ν*) for host genotypes ordered from the most resistant (G1) to the most permissive (G5) ones. Very interestingly, a significant negative regression coefficient exists for both traits [*AUDPS*: *−*1.313·10^−2^ ±0.383·10^−2^ (*R* = 0.459, *F*_1,44_ = 11.724, *P* = 0.001) and *I*: *−*2.576·10^−3^ ±0.790·10^−3^ (*R* = 0.4413, *F*_1,44_ = 10.621, *P* = 0.002)], suggesting that evolution always proceeded faster in the most restrictive hosts and slower in the most permissive ones.

**FIG. 6.**
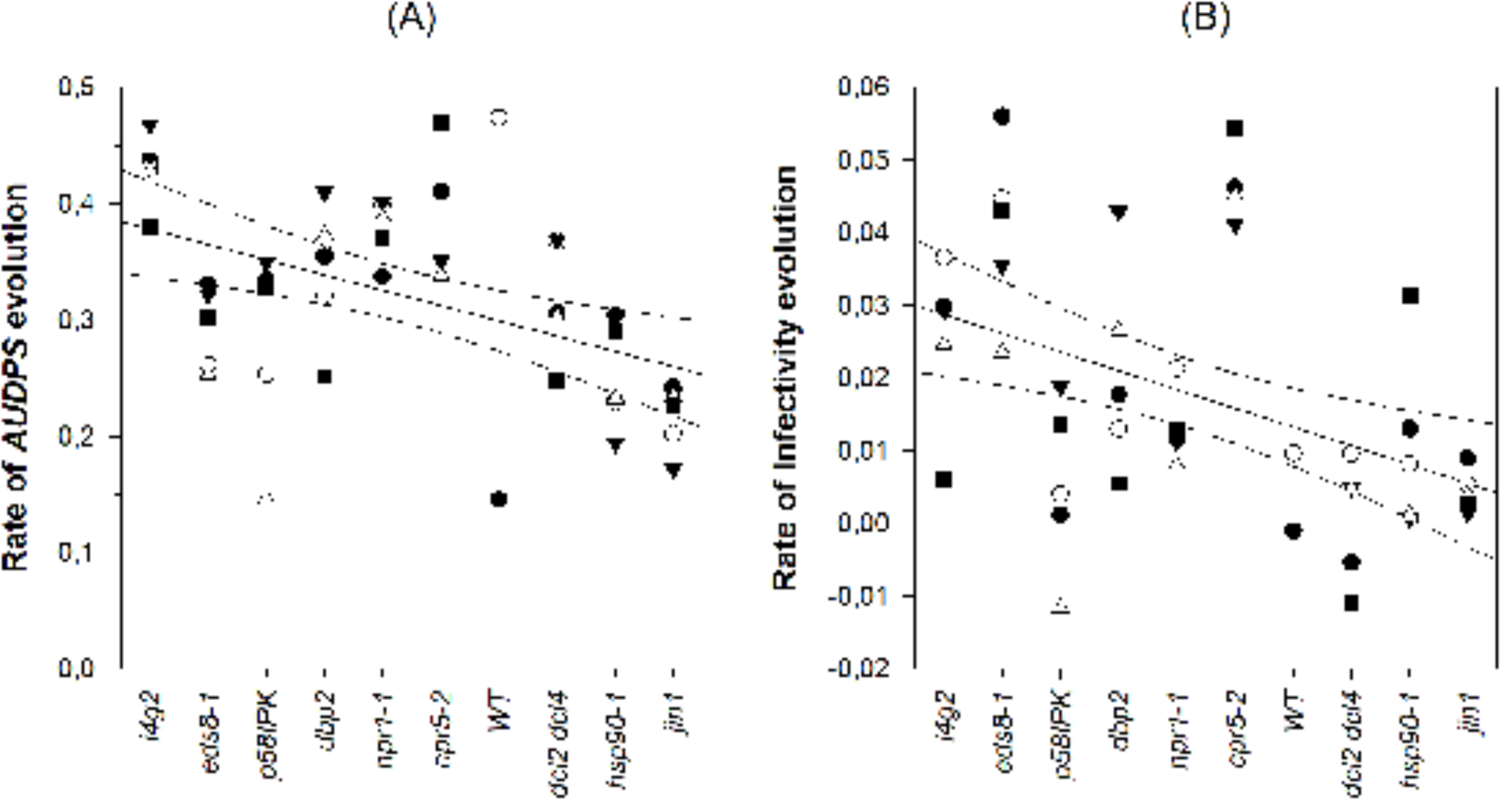
Estimated rates of evolution for *AUDPS* (A) and *I* (B) obtained from the fitting of an ARIMA(1,0,0) model (equation 2) to the data shown in figure 2 and figure 3, respectively. Different symbols represent different independent lineages evolved on the corresponding host genotype. Solid lines represent the linear regression of the data; dashed lines the 95% confidence intervals for the regression lines. In both cases, *A. thaliana* genotypes are ranked from the less to the more permissive to TuMV infection according to figure 1.

### Dissecting the role of specific defenses in TuMV evolution

The rates of TuMV phenotypic evolution estimated in the previous section were further analyzed in the context of the mutated defense signaling pathways or recessive resistances. Rates of evolution *ν_AUDPS_* and *ν_I_* were fitted to the MANOVA model shown in equation 3 of the Materials and Methods section. Figure 7 graphically summarizes the results of these analyses. First, we sought for differences in rates of evolution between permissive (soft selection) and restrictive (hard selection) hosts (figure 7A). The multivariate analysis showed that lineages evolved in the more restrictive hosts evolved significantly faster than those evolved in the more permissive hosts (*Λ* = 0.780, *F*_2,43_ = 6.080, *P* = 0.005). This effect was mostly driven by differences in the rate of infectivity evolution *ν_I_*, as shown by the corresponding univariate analyses, being on average 3.14 times faster in the most restrictive hosts. This observation confirms the results shown in the previous section.

**FIG. 7.**
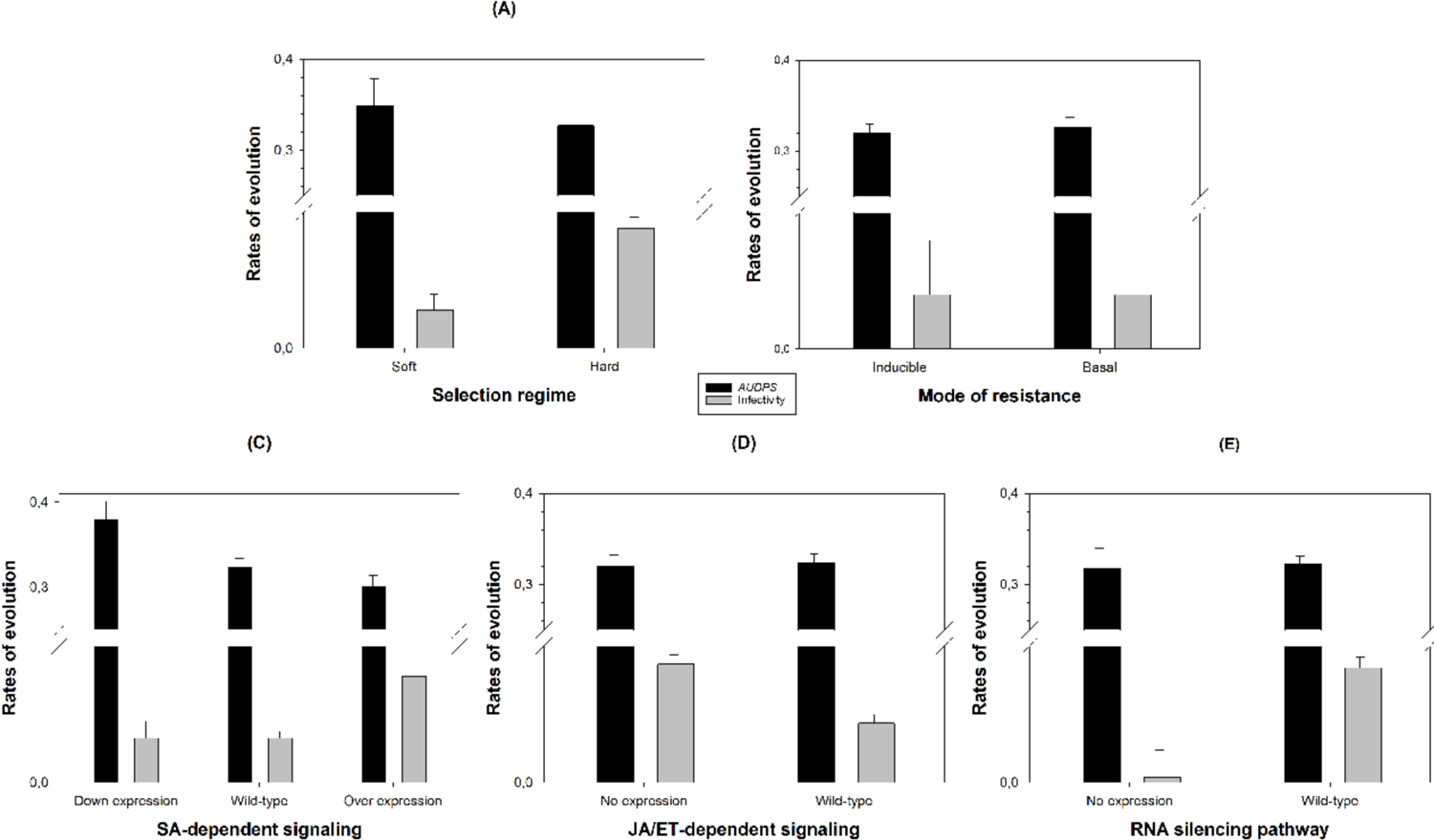
The rates of evolution for *AUDPS* and *I* differ among possible mechanisms of resistance to infection. Rates of evolution are expressed in log-scale to better visualize the slowest rates of *I* evolution. Data were fitted to the MANOVA model defined by equation. 3. Error bars represent ±1 SEM.

Next, we evaluated whether differences exist between the rates of phenotypic evolution for viruses evolved in host genotypes carrying mutations affecting basal or inducible resistances (figure 7B; WT data for this analysis excluded). In this case, the multivariate analysis found a small, yet significant effect (*Λ* = 0.790, *F*_2,34_ = 4.507, *P* = 0.018), though none of the two univariate analyses had enough power to detect it (indeed, changes went in opposite directions: 1.02-fold larger for *ν_AUDPS_* in the plants with mutations in basal resistances but 1.31 times larger for *ν_I_* in the plants with mutations in the inducible resistances).

Figure 7C shows the comparation of rates of TuMV phenotypic evolution for lineages evolved in plant genotype mutants for the SA-dependent signaling defense pathways. In this case we have three categories compared to WT: mutations inducing a down-expression of the pathway, mutations with no effect on this pathway and mutations resulting in overexpression. Again, the MANOVA found highly significant differences among the three categories (*Λ* = 0490, *F*_4,84_ = 8.996, *P* < 0.001). The univariate analyses showed that most of the differences were due to a 2.38 times faster evolution of *I* in the host genotypes over-expressing the SAR defenses, than in those with a WT-like or even down-expression of the SA signaling which results in a reduced SAR.

SA- and JA/ET-dependent pathways have been considered as antagonistic defenses (see Introduction), although more recent evidence points toward an active crosstalk between both (van Wees et al. 2000; Pieterse et al. 2012; Mine et al. 2017). Under the antagonistic hypothesis, we should expect the opposite trend among lineages evolved in plants with affected JA/ET signaling than the one observed for the lineages evolved in plants with affected SA signaling (figure 7C). Figure 7D shows the data of evolution rates classified according to whether host genotypes had mutations affecting the JA/ET defense signaling pathway that resulted ISR. The MANOVA found a significant difference between the two categories (*Λ* = 0.766, *F*_2,43_ = 6.580, *P* = 0.003), entirely driven by differences in *ν_I_*, as confirmed by the univariate analyses. Indeed, as expected, the rates of evolution for infectivity were 2-fold faster for TuMV lineages evolved in the ISR deficient hosts, in agreement with the SAR/ISR antagonistic hypothesis.

Finally, we tested the effect of knocking down the host’s RNA-silencing pathway in the rates of TuMV evolution (figure 7E). In this case, the multivariate analysis also found highly significant differences between rates of viral evolution in plants with a fully functional RNA-silencing pathway and the silencing-deficient *dcl2 dcl4* mutant (*Λ* = 0.850, *F*_2,43_ = 3.801, *P* = 0.030). The average *ν_I_* values were 21-fold larger in plants expressing a fully functional RNA-silencing pathway than in *dcl2 dcl4* plants unable of siRNA production.

In conclusion from this section, we found that regardless of the defense signaling pathway, viruses evolved in permissive plants always evolve slower than their counterparts evolved in more restrictive plant genotypes.

### Permissiveness of *A. thaliana* genotypes to infection drives the evolution of TuMV lineages into specialists or generalists

To analyze the specificity of adaptation of each evolved TuMV lineage, we performed a full cross-infection experiment. In this experiment, the 44 lineages evolved in the mutant genotypes (excluding the two WT-evolved lineages) were inoculated into ten plants of the nine *A. thaliana* mutant genotypes used in the evolution experiments (excluding WT plants). The presence of symptoms in each plant was recorded daily for up to 12 dpi. This dataset was analyzed using two different approaches, first an ANOVA-like that evaluates the effect of host genotypes, viral lineages and their interactions (Schmid-Hempel 2011); and second, the inference and characterization of nestedness, modularity and specialization of a bipartite infection network (Weitz et al. 2013). For the first approach we fitted the presence/absence of symptoms data to the logistic regression model shown in equation 4 of the Materials and Methods section. The results are shown in table 3. A test host genotype (*TG*) factor describes the effect of differences among host genotypes, irrespective of the infecting viral lineage; and thus, characterizes whether some host genotypes are more susceptible than others to infection. As shown in table 3, *TG* has a highly significant effect, both by itself and in the interaction with *t*. The magnitude of the net effect was very large (η^2^_P_ = 0.569), while its interaction with *t* was rather small in magnitude (η^2^_P_ = 0.010). On average, the most susceptible host genotypes were *jin1*, *dcl2 dcl4* and *cpr5-2* (f^-^ = 1.000), whereas the most resistant host genotypes were *p58^IPK^*(f^-^ = 0.459 ±0.013) and *eds8-1* (f^-^ = 0.233 ±0.013). Similarly, a highly significant local host genotype (*LG*) effect exists by itself (of large magnitude η^2^_P_= 0.288) as well as in combination with *t* (though the magnitude of the interaction was rather small in magnitude η^2^_P_ = 0.005), which means that the observation depends on the viral lineage independently of the host genotype. On average, the lineages evolved in *eds8-1* (f^-^ = 0.997 ±23.710) and *dbp2* (f^-^= 0.994 ±26.108) induced symptoms faster; while in contrast, lineages evolved in *i4g2* (f^-^ = 0.788 ±279.090) showed symptoms much slower. Finally, and most interestingly, a highly significant interaction term *TG×LG* (as well as in combination with *t*) indicates that the outcome depends on the particular combinations of host genotypes and TuMV lineages. Indeed, the magnitude of the interaction effect was large (η^2^_P_ = 0.314), though its dependence with dpi was not so relevant (η^2^_P_ = 0.048). For example, in the most susceptible genotypes, where *cpr5-2* perfectly illustrates this case: lineages evolved in *cpr5-2*, *dbp2*, *dcl2 dcl4*, *eds8-1*, and *p58^IPK^* all show f^-^ = 1.000, whereas lineages evolved in *i4g*2 and *jin1* show much lower f^-^ values (0.674 ±0.028 and 0.849 ±0.032, respectively). In the opposite extreme, *e.g*., *eds8-1*-evolved lineages show f^-^= 0.575 ±0.050 while *jin1*-evolved ones show f^-^= 0.124 ±0.030; while all other lineages show intermediate f^-^values.

**Table 3.** Results of the probit regression testing for the specificity of adaptation. The different model factors are defined in equation. 4.

| Source of variation | LRT | $df$ | $P$ | $\eta_p^2$ | $1 - \beta$ |
| --- | --- | --- | --- | --- | --- |
| $\phi$ | 21224.493 | 1 | $< 0.001$ | 0.959 | 1.000 |
| $t$ | 25995.056 | 1 | $< 0.001$ | 0.815 | 1.000 |
| $TG$ | 295.160 | 8 | $< 0.001$ | 0.569 | 1.000 |
| $t \times TG$ | 568.212 | 8 | $< 0.001$ | 0.010 | 0.981 |
| $LG$ | 20.545 | 8 | 0.008 | 0.288 | 0.661 |
| $t \times LG$ | 30.200 | 8 | $< 0.001$ | 0.005 | 0.804 |
| $L(LG)$ | 207.257 | 35 | $< 0.001$ | 0.385 | 1.000 |
| $t \times L(LG)$ | 235.073 | 35 | $< 0.001$ | 0.016 | 0.984 |
| $TG \times LG$ | 258.888 | 64 | $< 0.001$ | 0.314 | 1.000 |
| $t \times TG \times LG$ | 365.512 | 64 | $< 0.001$ | 0.018 | 0.961 |
| $TG \times L(LG)$ | 788.661 | 280 | $< 0.001$ | 0.048 | 0.999 |
| $t \times TG \times L(LG)$ | 1033.649 | 280 | $< 0.001$ | 0.042 | 0.993 |

LRT: likelihood ratio test distributed as a $\chi^2$ ; $\eta_p^2$ : magnitude of the effect; $1 - \beta$ : power of the test.

For the second analytical approach, we used the full cross-infection dataset to estimate the corresponding *AUDPS* values, thus creating an infection matrix of 44 rows [*L*(*LG*)] by nine columns (*TG*). This matrix was transformed into a new binary matrix in which a value of 1 meant that the *AUDPS* for a given TuMV lineage, in a particular *TG,* was greater or equal than the one observed for its *LG* and 0 otherwise. Figure 8A shows the packed infection matrix with rows and columns organized to better highlight its nestedness and modularity (Weitz et al. 2013). The matrix could be also represented as a bipartite infection network (figure 8B) that provides the same information but in a more graphical way. The infection network was significantly nested (*T* = 14.706, *P* < 0.001), with the most permissive genotypes *jin1* and *dcl2 dcl4* being susceptible to most viral lineages (universally susceptible hosts), while the most resistant genotype *eds8-1* and *p58^IPK^* were only successfully infected by some lineages that were precisely evolved in these two genotypes. Likewise, the most generalist viral lineages were *eds8-1*/L4, *p58^IPK^*/L3 and *p58^IPK^*/L4 that infected the nine mutant host genotypes (figure 8) with equal efficiency (universally virulent viruses). On another hand, the most specialized viral lineages that were able to successfully infect only their local hosts, were all surviving lineages evolved in *cpr5-2*, four lineages evolved in *dcl2 dcl4* and four lineages evolved in *jin1* (figure 8). Therefore, we conclude that local host’s resistance to infection positively correlates with the host range of the evolved lineages: more resistant host genotypes selected for very generalist viruses while less resistant host genotypes selected for very specialized viruses, in agreement with a GFG infection model (Weitz et al. 2013). Next, we computed Blüthgen et al. (2006) specialization indexes for the packed matrix (figure 8). Firstly, we evaluated the standardized species-level measure of partner diversity *d’*. *d’* ranged from zero for the most generalist TuMV evolved lineages (*eds8-1*/L4, *p58^IPK^*/L3 and *p58^IPK^*/L4) to one for the most specialist ones (the four surviving *cpr5-2*-evolved lineages, *dbp2*/L2, *dcl2 dcl4*/L2 - L5 and *jin1*/L1 - L4) (last column in figure 8A). Secondly, we computed the network-level specialization index $^#^, obtaining a value of $^#^= 0 (uncorrected *H*_2_ = 5.024, $^#^= 3.534, $^#^= 5.024; *P* < 0.001),which means that, overall, the binary infection network shows a degree of specialization. Most of the lineages were able of infect one or a few host genotypes, a result mainly driven by those lineages evolved in the less resistant host genotypes.

**FIG. 8.**
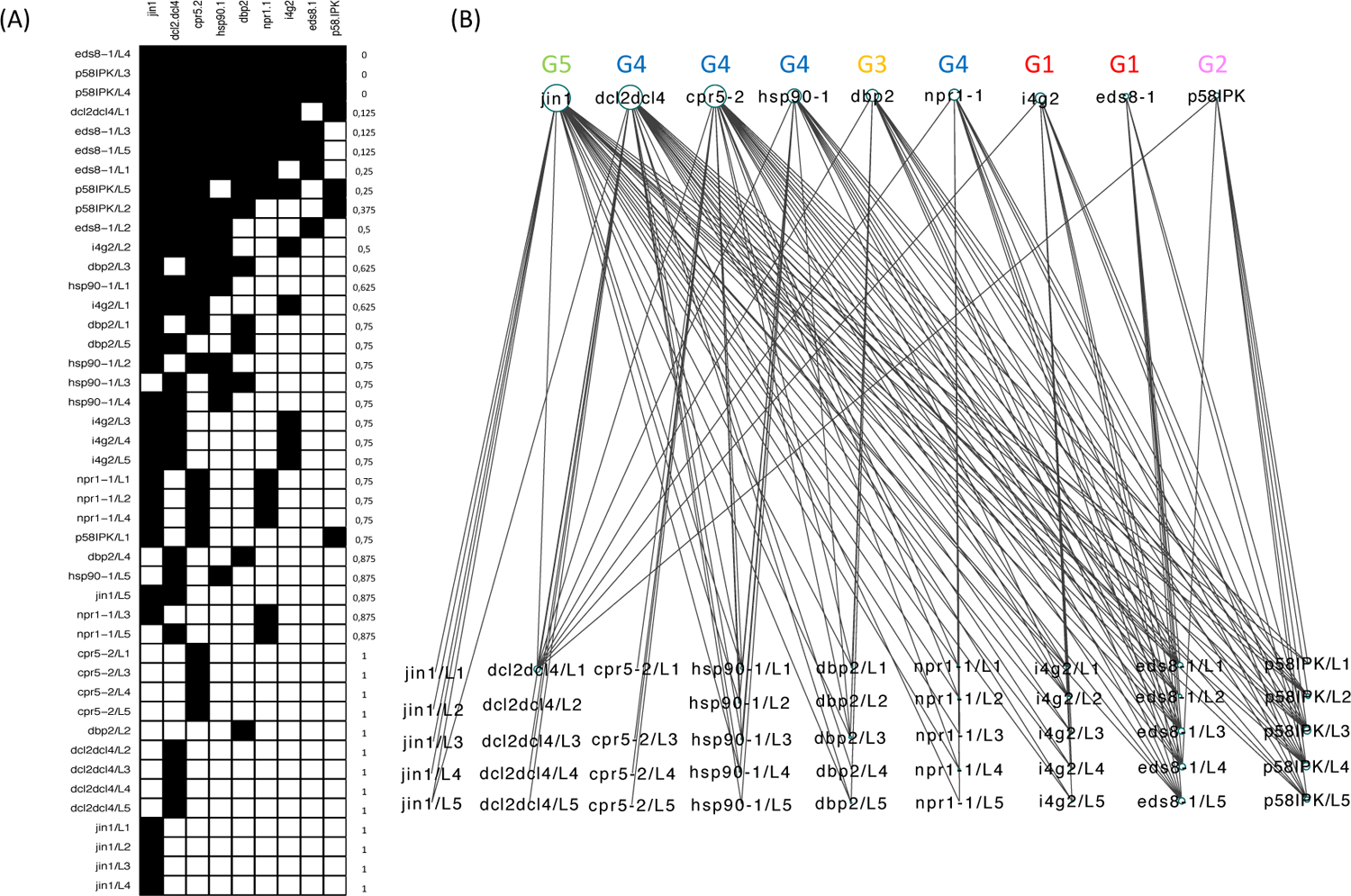
Analysis of the full cross-infection matrix. (A) Packed matrix that highlights its nested structure, compatible with a gene-for-gene infection model. Black squares represent cases in which the *AUDPS* values were equal or greater than those observed for the corresponding lineage in its local host. Last column shows the species-level specialization index *d’* (*d’* = 0 means most generalist pathogen, *d’* = 1 means most specialist pathogen). (B) Bipartite host genotype - viral lineage infection network. The size of the nodes is proportional to their degree. The phenotypic groups defined in figure 1 are indicated above the host genotype.

Finally, we sought to explore the modularity of the infection network. Indeed, from an ecological standpoint, a significant network-level degree of specialization means that some TuMV lineages do not interact well with some of the *A. thaliana* genotypes, thus creating the possibility of modularity. In this context, a module will refer to an aggregated set of viral lineages and their hosts characterized by more interactions within the module than between modules (Newman 2006; Dormann and Strauss 2014). To this end, we computed the *Q* modularity index, which ranges from zero, when the community has no more links within modules than expected by chance, to a maximum value of one. We found a small yet significant modularity (*Q* = 0.234, *P* = 0.047) in the infection matrix (figure 8A). What are the implications of modularity in TuMV pathogenesis? Modularity is expected for a MA infection model (Weitz et al. 2013). More similar host genotypes, *e.g*., those with mutations affecting the same signaling pathway, may be selecting for viral lineages with similar properties, thus being able of infect a subset of plant genotypes with equal efficiency. An alternative hypothesis that can explain modularity may be parallel evolution of lineages evolved into the same host genotype; the most representative cases of this possibility are the lineages evolved in *cpr5-2*, *dcl2 dcl4* and *jin1*. To distinguish between these two hypotheses, we computed a reduced matrix averaging the observed *AUDPS* among lineages evolved in the same host genotype and transforming this nine-by-nine infection matrix into its corresponding packed binary matrix, as described above.

If the reduced matrix still shows significant modularity, the first hypothesis will hold on, but if modularity vanishes by averaging lineages evolved in the same host genotype, then the alternative hypothesis will be more parsimonious. The reduced matrix is still significantly nested (*T* = 13.064, *P* < 0.001), but modularity was not significant anymore (*Q* = 0.286, *P* = 0.137). This supports the hypothesis that the observed modularity was driven by convergent phenotypic evolution of lineages evolved into the same host genotype rather than by the overlap of selective pressures by genotypes carrying mutations in the same defense mechanism.

### Genomic differences/similarities among TuMV lineages and VPg as a potential target of selection

Finally, we have explored the molecular changes experienced by the TuMV evolved lineages. Full genomic consensus sequences were determined for 45 evolved lineages. A total of 119 mutational events have been observed, affecting 73 nucleotide positions (supplementary figure S3A and table S1, Supplementary Material online). According to the type of nucleotide substitution involved, 98 were transitions and 21 transversions; regarding their effect on the protein sequence, 30 were synonymous and 89 nonsynonymous. Interestingly, some mutations, including 13 nonsynonymous and four synonymous, have been observed multiple times in independent lineages. As discussed in the Materials and Methods section, treating each lineage as an observation and each host genotype as a subpopulation, the average nucleotide diversity within host genotypes, referred only to the 73 polymorphic sites, was *π_S_* = 0.067 ±0.007. On the other hand, the nucleotide diversity for the entire sample was *π_T_* = 0.073 ±0.008. Hence, the estimated interhost genotypes nucleotide diversity was *δ_ST_* = 0.006 ±0.003, which results in a coefficient of nucleotide differentiation (Nei 1982) of *N_ST_* = 0.081 ±0.039, a value significantly greater than zero (*z* = 2.077, *P* = 0.019). Thus, we conclude that minor yet significant genetic differentiation has been generated among viral lineages replicating in different host genotypes. To assess whether selection played a role in virus genetic differentiation among *A. thaliana* genotypes, we performed a Tajima’s *D* test (Tajima 1989) and found that it was significantly negative (*D* = *−*2.496, *P* = 0.006).

Next, we sought to characterize the distribution of mutational events along the nine non-overlapping cistrons (supplementary figure S3B, Supplementary Material online). The frequency of mutations per cistron, relative to the length of the corresponding cistron, was fitted to the logistic regression model shown in equation 5 of the Materials and Methods. Highly significant differences exist (*χ*^2^ = 143.206, 9 d.f., *P* < 0.001), yet entirely due to the ∼11-fold larger mutation frequency observed in the *VPg* cistron relative to the rest of the genome (supplementary figure S3B, Supplementary Material online). Notice that all mutations observed in *VPg* are nonsynonymous and that all lineages except *eds8-1*/L3 carry at least one mutation in this cistron (supplementary figure S3A and table S1, Supplementary Material online).

Convergent nonsynonymous mutations are, *a priori*, good candidates for adaptive mutations. Mutation CP/V148I appears in two lineages evolved in *cpr5-2* (supplementary figure S3A and supplementary table S1, Supplementary Material online), mutation CP/S70N appear in lineage *dcl2 dcl4*/L1 and three lineages evolved in *jin1* (supplementary figure S3A and table S1, Supplementary Material online), and two different mutations affecting CP amino acid 112, CP/D112G (lineage *npr1-1*/L4) and CP/D112A (lineage *i4g2*/L5), that result in a similar replacement of side chains. In addition, 10 nonsynonymous mutations in *VPg* are shared by several lineages (supplementary figure S3A and table S1, Supplementary Material online). Out of these 10 cases, three seem particularly promising candidates. They all are located in a narrow region of VPg (residues 113, 115 and 118) (supplementary figure S3A and table S1, Supplementary Material online). Firstly, mutations G6237A, G6237C, A6238G, and U6239A all affect the same codon, resulting in amino acid replacements D113N, D113H, D113G, and D113E, respectively. The expected functional effect of these mutations was evaluated using the algorithms implemented in SNAP2 (Hecht, Bromberg and Rost 2015). D113E is predicted to be functionally neutral (score = *−*76, accuracy = 87%), while D113G (32, 66%), D113H (43, 71%) and D113N (30, 66%) are predicted to have a functional effect. Among these three amino acid replacements, D113G, which shows an intermediate value of the functional effect score, is particularly interesting, involving a very strong change in the side radical from a long negatively charged one to a small nonpolar one. This replacement has been observed in lineages *cpr5-2*/L3, *cpr5-2*/L4, *cpr5-*2/L5, *npr1-1*/L1, *npr1-1*/L5, *hsp90-1*/L2, and WT/P. All four mutant hosts where these lineages evolved belong to the phenogroup G4 (figure 1). Secondly, mutations A6243C, A6243G and A6244G also affect the same codon, resulting in amino acid replacements N115H (*−*42, 72%), N115E (*−*66, 82%) and N115S (*−*30, 61%). All three are predicted to be functionally neutral. Thirdly, mutations U6252 and G6253A affect the same codon and result in amino acid replacements R118C (*−*10, 53%) and R118H (56, 78%), respectively. The most frequent among these two replacements is R118H, which represents a conservative change among large positively charged side radicals, yet has a strong expected functional effect. R118H has been observed in four out of five TuMV lineages evolved in *jin1* (phenotypic group G5 in figure 1). D113G thus represents a case of convergent evolution non-specific of the local host genotype (*i.e*., a candidate for a generalist mutation) while R118H represents a mutation highly specific of the local host genotype (*i.e.*, a specialist mutation).

To further characterize the possible adaptive value of mutations VPg/D113G and VPg/R118H, we created by site-directed mutagenesis the two mutated versions of *VPg* and cloned them into the ancestral p35STunos infectious clone (Chen et al. 2003). Viruses were recovered from these clones and three disease-related phenotypic traits (*e.g*., *AUDPS*, *I* and *SS*) evaluated both in the WT plants as well as in their corresponding local hosts (*i.e*., *cpr5-2* for mutant VPg/D113G and *jin1* for mutant VPg/R118H). Figure 9 summarizes the results of these experiments. Regarding VPg/D113G (figure 9A - C), the results strongly depend on the combination of phenotypic trait and plant genotype: it shows a significant negative effect in both *AUDPS* ( *−*14.2%; *z* = 1.687, *P* = 0.046) and *I* (*−*23.9%; *z* = 6.344, *P* < 0.001) but a positive one in *SS* (16.7%; *z* = 2.597, *P* = 0.005) when evaluated in WT plants. However, it shows a largely negative yet not significant effect in *cpr5-2* plants for *AUDPS* (*−*60.8%; *z* = 1.055, *P* = 0.146) and significant negative effects both for *I* (*−*60.1%; *z* = 10.614, *P* < 0.001) and *SS* (*−*46.7%; *z* = 3.228, *P* = 0.001). These negative effects on the local host genotype *cpr5-2* do not support a possible beneficial effect of this mutation by itself. Regarding the possible adaptive effect of the amino acid replacement VPg/R118H (figure 9D - F), the results are also dependent on the combination of phenotypic trait and host plant genotype. When tested in WT plants, the mutation has significant negative effects both for *AUDPS* (*−*11.4%; *z* = 2.29, *P* = 0.013) and *I* (*−*9.0%; *z* = 2.749, *P* = 0.003) but a significantly positive effect for *SS* (43.6%; *z* = 7.881, *P* < 0.001). In contrast, in the local host *jin1* the effect was positive in all three traits; significant for *AUDPS* (9.2%; *z* = 1.683, *P* = 0.046) and *SS* (68.2%; *z* = 12.447, *P* < 0.001) but not for *I* (0.3%; *z* = 1.342, *P* = 0.090). These results support the conclusion that the amino acid replacement VPg/R118H has an overall beneficial effect on the local host genotype *jin1* in which it was selected.

**Fig. 9.**
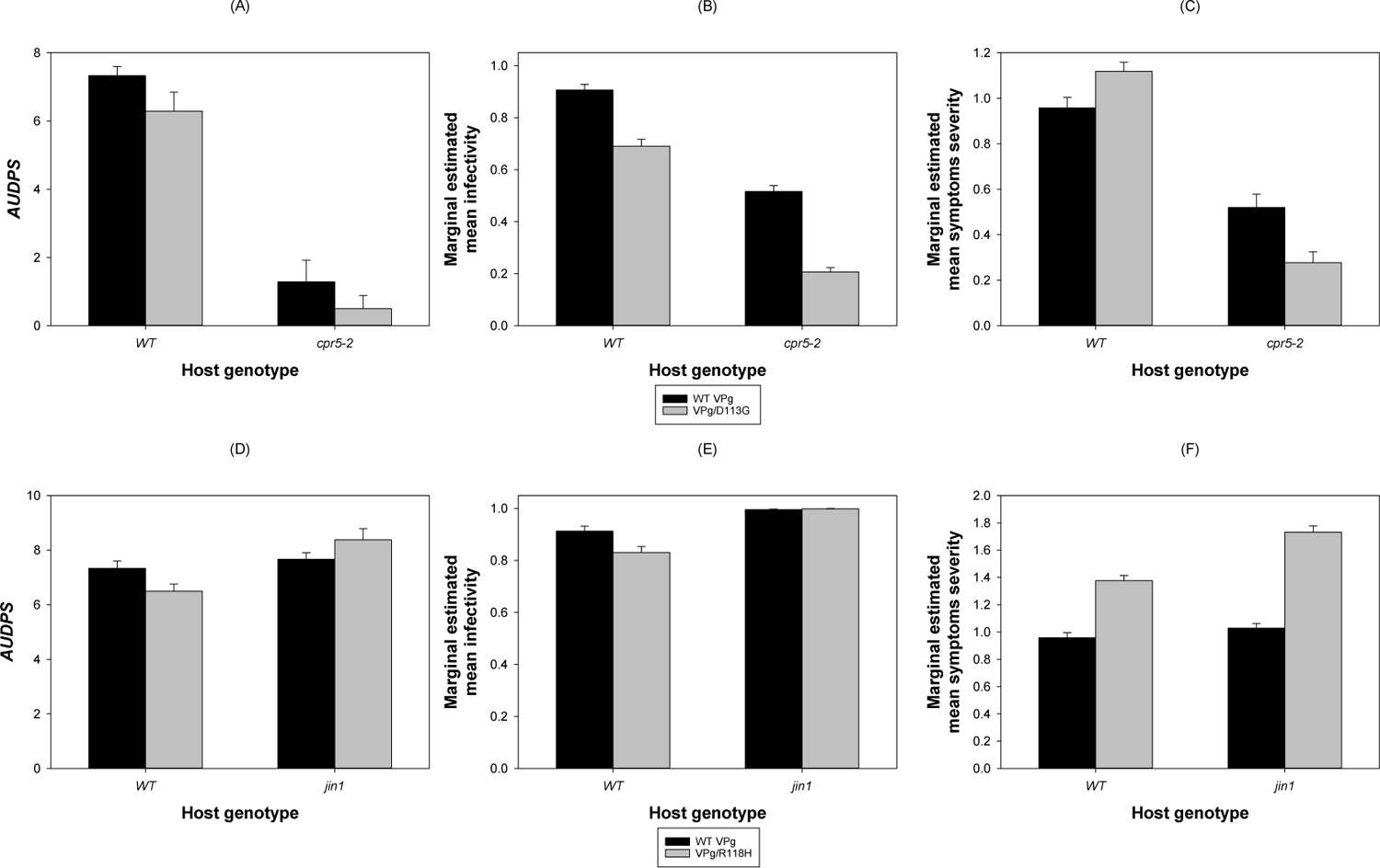
Phenotypic disease-related traits (*AUDPS*, *I* and *SS*) of TuMV VPg/D113G relative to WT VPg. (A) – (C) Evaluated in the WT *A. thaliana* and the *cpr5-2* mutant genotype. (D) – (F) Evaluated in the WT *A. thaliana* and the *jin1* mutant genotype. Error bars represent ±1 SEM.

## Discussion

### Interaction between TuMV and different plant defenses

In this study we have explored the effect that mutations in different host disease signaling pathways or in recessive resistance *S* genes have in the outcome of virus evolution. Our experimental pathosystem consisted of two well studied components: the model plant *A. thaliana* and one virus with high prevalence in natural populations on this host, TuMV (Pagán et al. 2010). Among the disease signaling pathways, we have studied genes involved in the SA, JA/ET and RNA-silencing; among the *S* genes, we have included heat shock proteins, transcription factors and components of the translation machinery. In a preliminary set of experiments, we compared phenotypic responses of different plant mutant genotypes to TuMV infection. Our results were, in some cases, at odds with the expected phenotypes for *R* genes, the most unexpected one being the result for *jin1*. *JASMONATE INSENSITIVE 1* (*JIN1*) is a negative regulator of SA-mediated defense responses, henceforth the mutant *jin1* has a constitutive expression of SAR (Laurie-Berry et al. 2006). Surprisingly, it turns out to be the most sensitive genotype to TuMV infection, showing enhanced symptoms of disease (phenogroup G5). This may reflect the fact that the function of genes involved in disease signaling have been mostly defined in terms of plant interactions with biotrophic and necrotrophic bacteria and fungi but rarely in response to virus infections. Mutant genotypes only affecting components of the ISR pathway (*e.g*., *coi1-4*) behaved as WT plants in response to TuMV infection, thus confirming previous reports that ISR was inefficient against viral infections (Ton et al. 2002; Loebenstein 2009; Pieterse et al. 2009). However, mutant *eds8-1*, which avoids ISR but enhances SAR (Love et al. 2007) turned out to be among the most resistant genotypes to infection (phenogroup G1). In agreement with our findings of a variable response to SA-signaling, Singh et al. (2004) discussed examples of viruses in which SA-dependent responses to viral infection were strongly dependent on the virus used in the experiments. Replication of viruses such as alfalfa mosaic virus, potato virus X or turnip vein clearing virus was inhibited by treating plants with SA, while cucumber mosaic virus appeared unaffected by the treatment and accumulated to normal levels.

The RNA-silencing pathway is considered as the main plant defense against viral pathogens (Voinnet 2001), with *DICER LIKE 2* (*DCL2*) and *DICER LIKE 4* (*DCL4*) encoding for the two dicer enzymes responsible for generating the 22- and 21-nucleotides long antiviral siRNAs. Therefore, the double mutant *dcl2 dcl4* was expected to show a highly sensitive response to TuMV infection. However, infected plants were, overall, hardly distinguishable from the WT plants in their response to infection (supplementary figures S2 and S3, Supplementary Material online). A possible explanation would be the strong suppressor activity of HC-Pro in the WT plants that effectively counteracts the defense mechanisms (Kaschau and Carrington 2001). Indeed, it has been shown that the outcome of the interplay between plant RNA-silencing response and potyviruses is mostly driven by the efficiency of the viral HC-Pro suppression activity (Lin et al. 2007; Torres-Barceló et al. 2008).

The response to infection of genotypes carrying mutations in *S* genes was, in general, consistent with the *a priori* expectation. Mutant *i4g2* belonging to phenogroup G1, that is defective for the eIF4(iso)E factor, shows a strong resistance to TuMV infection (Nicaise et al. 2007; Charron et al. 2008). Likewise, in agreement with previous descriptions, mutant *dbp2* belonging to phenogroup G3, shows enhanced resistance to potyvirus infection (Castelló et al. 2011). Finally, mutant *hsp90-1* turns out to show a response to TuMV infection equivalent to the WT plants, likely due to its functional redundancy with other heat-shock proteins present in the cell that may be coopted by the virus to assist in their expected functions (Verchot 2012).

An interesting finding was the effect of the knocked down expression of gene *P58^IPK^* on TuMV replication. In mammals P58^IPK^, a tetratricopeptide repeat (TPR)-containing protein, is recruited by viruses (*e.g*., influenza A virus) to inhibit interferon activation and cell death mediated by the dsRNA-activated protein kinase (PKR), thus favoring viral spread (Goodman et al. 2011). In our experiments, *p58^IPK^* mutant plants showed enhanced tolerance to infection by showing weaker symptoms than WT plants despite the same level of virus accumulation (supplementary figure S1, Supplementary Material online). In sharp contrast, Bilgin et al. (2003) found that infection of *p58^IPK^*-silenced *Nicotiana benthamiana* and *A. thaliana* plants infected with tobacco mosaic virus led to death. Given the limited information available from other plant-virus pathosystems, additional experiments would be necessary to shed light into the potential antiviral role in plants of this gene.

An important conclusion of our study is that the generally accepted model of plant defense signaling pathways, mainly based in experiments done with bacteria and fungi, may not describe well the interaction between plants and viruses. By contrast, recessive *S* gene-based resistance seems to better explain differences in susceptibility to infection and viral accumulation, thus being a more promising target for future development of resistant plants.

### Evolution of specialist and generalist viral strategies depends on the host genotype

The GFG and MA models of host-virus interaction modes represent the two ends of a continuum of possible outcomes (Agrawal and Lively 2002). The most relevant difference between both models regards the expected genetic heterogeneity in both host and virus populations. With a pure GFG interaction, the susceptible host types are expected to disappear and the resistant types to dominate the population. *Vice versa*, the most virulent virus allele would become fixed in the population at the cost of mild alleles. However, constitutive activation of defenses is known to be costly for *A. thaliana* [*e.g*., SA-related defense responses pay a fitness cost in absence of pathogens (Traw et al. 2007)] and high virulence usually comes with a cost in terms of pathogen’s transmission (Acevedo et al. 2019). Hence, a pure GFG strategy seems unlikely to be achieved. By contrast, with a pure MA interaction, negative frequency-dependent selection emerges, such that rare *A. thaliana* resistance alleles have advantage and, as a result, a genetic polymorphism shall be maintained (Schmid-Hempel 2011). Natural populations of *A. thaliana* contain considerable amount of genetic variability for tolerance (Pagán et al. 2008) and for immunity-related genes (Todesco et al. 2010, van de Weyer et al. 2019), thus suggesting that an arms race between pathogens and plants is still ongoing but it is unclear whether it may result in pure GFG or MA interactions or lie somewhere in between. Evolution experiments with such different pathosystems as TuMV-*A. thaliana* (González et al. 2019), *Octosporea bayeri*-*Daphnia magna* (Altermatt and Ebert 2008) and *Serratia marcescens*-*Caenorhabditis elegans* (Gibson et al. 2020; White et al. 2020) have produced congruent results: parasites exposed to heterogeneous host populations evolved significantly lower virulence than parasites exposed to homogeneous host populations. However, a significant difference exists among pathosystems: while viruses exposed to genetically heterogenous host populations evolved as no-cost generalists, evolution of generalism in more complex parasites was constrained by a fitness tradeoff, as expected for the jack-of-all trades hypothesis (Bedhomme et al. 2015).

Our results, as well as those by Hillung et al. (2014) and González et al. (2019) have shown the evolution of significantly nested bipartite infection networks, a finding compatible with the existence of a combination of specialist and generalist viruses and of more permissive and resistant host genotypes. Indeed, these studies have also shown that more permissive hosts selected for more specialized viruses while more resistant hosts selected for more generalist viruses, here matching the predictions of the GFG model. Our observation of small yet significant modularity in the infection network was easily explained by convergent evolution of TuMV lineages evolved in the same host genotype. However, it has been recently shown in a long-term survey of the prevalence of different plant viruses in different hosts and habitats that nestedness and modularity in host-pathogen infection networks is possible due to the spatially patched distribution of habitats and temporal successions of plant species (Valverde et al. 2020): small spatial scales create modularity that coexist with global nestedness. This pattern may change spatially and temporally but remains stable over long evolutionary timescales.

### Role of natural selection

We found evidence of significant genetic differentiation among TuMV lineages evolved in different plant genotypes. To test whether these differences were driven by selection we performed a Tajima’s *D* test (Tajima 1989). The resulting negative *D* value was significant, which is compatible with the action of purifying selection, the presence of slightly deleterious mutations segregating in the populations or fast population expansions (Yang 2006). How to distinguish between these explanations? In independent fast expanding populations, many new mutations may be generated and rising in frequency in each population, thus being observed as singletons, mutations present in only one of the many coexisting genomes in each evolving TuMV lineage. Singletons inflate the number of segregating sites and thus cause *D* < 0. Indeed, this is the case here: 56 out of the 73 observed variable sites are singletons, thus the observed pattern of molecular diversity among lineages evolved in the same host genotype and in different host genotypes is likely to be due to the fast expansions of viral populations.

However, we have found additional evidence supporting the action of positive selection: the existence of a number of convergent nonsynonymous mutations arising in independent lineages. Some of these nonsynonymous mutations evolved in the same host genotype but others arise in different host genotypes (supplementary table S1 and figure S3, Supplementary Material online). Yet without a clear association with the particular signaling pathway, mutated *S* gene (table 1) or the phenogroup they belong to (figure 1). Interestingly, most of these convergent mutations happened in the *VPg* cistron, which also turns out to be the most variable one. VPg plays many essential roles in genome transcription (it is linked to the 5’-end of the viral genome and provides the hydroxyl group that primes the synthesis of the complementary strains by the viral RdRp), translation [directly interacts with the eukaryotic initiation factors eIF(iso)4E and eIF(iso)4G] and interacts with all other viral proteins (Bosque et al. 2014) and some of the host cell proteins (Martínez et al. 2016; Martínez et al. 2020). Indeed, in previous evolution experiments with potyviruses, *VPg* has also been shown to be an important target of selection. For example, Agudelo-Romero et al. (2008) found that a single amino acid replacement in VPg was enough to largely increase TEV *I*, *SS*, *VL* and in *A. thaliana*. Similarly, Gallois et al. (2010) found that *A. thaliana* plants with knock-out mutations in the *eIF(iso)4E*, *eIF(iso)4G1* and *eIF(iso)4G2* genes were resistant to TuMV infection. Two mutations in the VPg (E116Q and N163Y) were enough to overcome this resistance and return to the original infection phenotype, though yeast-two hybrid assays showed that none of these mutations affected the binding of VPg with eIF(iso)4E (Gallois et al. 2010). As a final example, one of the most extensively used resistance genes against potato virus Y in commercial pepper cultivars is *pvr^2^*, which has many different alleles (Nicaise et al. 2007; Charron et al. 2008). The *pvr^2^* locus encodes for the eIF4E factor which, as mentioned above, physically interacts with VPg. Interestingly, all the resistance-breaking viral isolates found so far contain mutations in the *VPg* cistron (Duprat et al. 2002; Moury et al. 2004; Ayme et al. 2006).

One of the two mutations identified by Gallois et al. (2010), VPg/E116Q, affects the same protein domain as mutations D113G and R118H identified and characterized here. The mechanism by which these two mutations may confer a selective advantage to TuMV lineages cannot be inferred from our studies. We found VPg/D113G in several lineages evolved in different host genotypes, while VPg/R118H was only found in the *jin1* lineage. Our assays failed to find a beneficial effect of VPg/D113G but confirmed the host-specific beneficial effect of VPg/R118H in *jin1* plants.

Unfortunately, we have not been able to exhaustively test all observed mutations and, hence, several others may still be potential candidates for adaptive mutations. More specifically, we have not tested possible epistatic interactions among mutations fixed in the same genome. For example, three lineages carry more than one nonsynonymous mutation in *VPg* (*dbp2*/L1 D113N and N115E, *eds8-1*/L1 N115H and E116G, and *eds8-1*/L4 H33Y, D113N and K121E) (supplementary table S1, Supplementary Material online). In addition, 25 lineages carry additional nonsynonymous mutations in at least one other cistron besides *VPg* (supplementary table S1, Supplementary Material online). Two lineages have up to four additional nonsynonymous mutations: (*i*) *cpr5-2*/L5 that has mutations in proteins P3/T318M-T326S, CI/V148I and VPg/D113G and (*ii*) *dcl2 dcl4*/L1 that has mutations in proteins P1/V54A, CI/I378V, VPg/R118C, and CP/S70N. Therefore, plenty of opportunities for epistatic effects both within and among cistrons exist, and will be evaluated in future experiments.

## Methods

### Plants, virus and growth conditions

A collection of 21 different *A. thaliana* mutants of the Col-0 accession were used for this study (table 1). In all experiments described below, plants were all maintained in a BSL2 climatic chamber under a photoperiod of 8 h light (LED tubes at PAR 90 - 100 µmol/m^2^/s) at 24 °C and 16 h dark at 20 °C and 40% relative humidity.

Prior to the inoculation experiments, we created a large stock of TuMV infectious saps. Saps were obtained from TuMV-infected *N. benthamiana* Domin plants inoculated with the infectious plasmid p35STunos that contains a cDNA of TuMV genome (GeneBank accession AF530055.2) under the control of the cauliflower mosaic virus 35S promoter and the *nos* terminator (Chen et al. 2003) as described elsewhere (González et al. 2019; Corrêa et al. 2020). This TuMV sequence variant corresponds to the YC5 isolate from calla lily (*Zantesdeschia* sp.) (Chen et al. 2003). After the plants showed symptoms of infection, they were pooled and frozen with liquid N_2_. This frozen plant tissue was homogenized into a fine powder using a Mixer Mill MM400 (Retsch GmbH, Haan, Germany). For inoculations, the 0.01 g of powder was diluted in 1 mL inoculation buffer (50 mM phosphate buffer pH 7.0, 3% PEG6000, 10% Carborundum) and 5 μL of the inoculum was gently rubbed into the plant leaves. Plants were all inoculated when they reached the growth stage 3.5 in the Boyes et al. (2001) scale. This synchronization ensures that all hosts were at the same phenological stage when inoculated.

### Phenotyping infected plants

Five different disease-related traits were measured for each infected plant for 18 dpi. (*i*) Change in dry weight (*ΔDW*) of the aerial part of infected plants, with a precision of 10 mg, relative to the corresponding noninfected controls (supplementary figure S1A; Supplementary Material online). (*ii*) Severity of symptoms (*SS*) evaluated in a semi-quantitative discrete scale (figure 1B in Corrêa et al. 2020) ranging from zero for asymptomatic infections to four for plants showing a generalized necrosis and wilting (supplementary figure S1B, Supplementary Material online). (*iii*) The area under the disease progress stairs (*AUDPS*), that summarizes the speed at which the disease progresses in a group of plants (supplementary figure S1C, Supplementary Material online) and takes values in the range (0, *N*) where *N* is the total number of plants included in the assay (Simko and Piepho 2012). (*iv*) Infectivity measured as the number of symptomatic plants out of the number of inoculated plants at 18 dpi (*I*; supplementary figure S1D, Supplementary Material online). And (*v*) viral load (*VL*) measured by absolute RT-qPCR as the number of viral genomes per ng of total RNA in the plant as described below (supplementary figure S1E, Supplementary Material online).

The five traits were not all independent but showed some significant pairwise positive correlations: *AUDPS* with *I* (*r* = 0.772, 19 df, *P* < 0.001), *SS* (*r* = 0.807, 19 df, *P* < 0.001) and *VL* (*r* = 0.472, 19 df, *P* = 0.031), and *I* with *SS* (*r* = 0.746, 19 df, *P* < 0.001).

Basically, *ΔDW* is orthogonal with the other four traits while the other four traits showed some degree of association. Interesting is the case of *VL*, which was only (weakly) correlated to *AUDPS*.

### Experimental evolution

Five TuMV lineages were evolved during 12 consecutive serial passages in each one of the nine selected mutant genotypes. In addition, two lineages were also evolved in WT plants. To begin the evolution experiment, ten 21-days old *A. thaliana* plants per lineage and genotype were inoculated as described above with the virus stock previously created from infected tissue of *N. benthamiana* plants inoculated with an TuMV isolate YC5 infectious clone. The next passages were made by harvesting the symptomatic plants at 14 dpi, preparing the infectious sap as described above and inoculating it to a new healthy population of ten plants. These saps were 1/10 diluted with inoculation buffer and used to inoculate the next batch of plants.

### Total RNA extractions

Tissue from each pool of infected plants per lineage, genotype and serial passage was collected, frozen with liquid N_2_ and preserved at *−*80 °C until it was homogenized into fine powder using a Mixer Mill MM400. Next, an aliquot of approximately 100 mg of grounded tissue was used for total RNA (RNAt) extraction. The RNAt was extracted with the Agilent Plant RNA isolation Mini kit (Agilent Technologies, Santa Clara CA, USA). Aliquots of RNAt per each sample were separated and their concentration adjusted at approximately 50 ng/μL to estimate viral accumulation by RT-qPCR (see below).

### Quantification of *VL*

*VL* of each plant sample per lineage, genotype and selected passage was quantified by absolute real-time quantitative RT-PCR (RT-qPCR) using standard curves and the primers TuMV F117 forward (5’-CAATACGTGCGAGAGAAGCACAC-3’) and F118 reverse (5’-TAACCCCTTAACGCCAAGTAAG-3’) that amplify a 173 nucleotides fragment from the *CP* cistron of TuMV genome, as previously described (Corrêa et al. 2020). Briefly, standard curves were constructed using ten serial dilutions of the TuMV genome, that was synthesized by *in vitro* transcription as detailed previously (Cervera et al. 2018), in RNAt extract obtained from healthy *A. thaliana* plants used as control in the experiments. Amplification reactions were run in a 20 μL volume using the GoTaq 1-Step RT-qPCR System (Promega, Madison WI, USA) and the recommended manufacturer’s instructions as described in Cervera et al. (2018) in an ABI StepOne Plus Real-time PCR System (Applied Biosystems, Foster City CA, USA). The cycling conditions consisted in: an RT phase of 5 min at 42 °C and 10 min at 95 °C followed by a PCR stage consisting in 40 cycles of 5 s at 95 °C and 34 s at 60 °C; and the final melt curve profile that consisted in 15 s at 95 °C, 1 min at 60°C and 15 s at 95 °C. Negative controls consisted of healthy RNAt plant extract (mock-inoculated non infected control) and water. For each sample, three technical replicates were quantified. The results were analysed using the StepOne software 2.2.2 (Applied Biosystems).

### TuMV genome amplifications and sequencing

Evolved viral genomes of the passage 12 from each lineage and genotype were amplified by high-fidelity RT-PCR using the AccuScript Hi-Fi (Agilent Technologies) reverse transcriptase and Phusion DNA polymerase (Thermo Scientific, Waltham MA, USA) following the manufacturer’s instructions. Each complete TuMV genome was amplified into three overlapping amplicons of 3114 (5’ fragment R1), 3697 (central region R2) and 3287 nucleotides (3’ fragment R3) using three primer sets. For RT reactions an aliquot of the corresponding RNAt (150-300 ng) was mixed with 0.25 μM of the 1R-P3 (5’-CGAGTAGTATCTTATAGCACAGCGCTCCGACC-3’), 2R-NIa (5’-TGTCTGGAATCGGTAGCAAATGTAGCTGAGTTGTG-3’) or 3R-polyAR (5’-TTTTTTTTTTTTTTTTTTTTGTCCCTTGCATCATATCAAATG-3’) primer to synthesize the R1, R2 or R3 cDNA fragment, respectively, that were denatured 5 min at 65 °C and cooled on ice. Then a mix containing AccuScript Hi-Fi 1*×* Buffer, 1 mM of dNTPs, 8 mM of DTT, 4U of Ribolock RNase inhibitor (Thermo Scientific) and 0.5 μL of AccuScript Hi-Fi (Agilent Technologies) was added up to a 10 μL volume. RT reactions consisted of 90 min at 42 °C to synthesize the cDNA followed by an incubation of 5 min at 70 °C to inactivate the enzyme. PCR reactions were performed in a 50 μL volume containing a mix of 1*×* Phusion Buffer, 0.4 μM of dNTPs, 0.2 μM of each primer, 0.5-1 μL of DMSO, 2U of Phusion DNA polymerase (Thermo Scientific) and 1 μL of the corresponding RT reaction. R1 fragment was amplified using the primer set 1F-5UTR (5’-GCAAACGCAGACCTTTCGAAGCACTCAAGC-3’) and 1R-P3 and the following PCR conditions: an initial denaturation of 30 s at 98 °C, 3 cycles of 10 s at 98 °C, 20 s at 67 °C and 2 min at 72 °C, 3 cycles of 10 s at 98 °C, 20 s at 65 °C and 2 min at 72 °C, and 32 cycles of 10 s at 98 °C, 20 s at 63 °C and 2 min at 72 °C, followed by a final extension step of 5 min at 72 °C. Fragments R2 and R3 were amplified with primer set 2F-P3 (5’-TGGGAGCTTGCGGATGGTGGATACACAATTC-3’) and 2R-NIa or 3F-NIa (5’-CTCGTTATATGGAGTCGGTTTCGGACCACTCATAT-3’) and 3R-polyAR, respectively and a PCR with the same denaturation and extension steps than the fragment R1 but different amplification steps: a stage consisting of 15 cycles of 10 s at 98 °C, 20 s at 67 °C and 2 min at 72 °C followed by 23 cycles of 10 s at 98 °C, 20 s at 65 °C and 2 min at 72 °C for fragment R2. For fragment R3 15 cycles of 10 s at 98 °C, 20 s at 67 °C and 2 min at 72 °C followed by 23 cycles of 10 s at 98 °C, 20 s at 65 °C and 2 min at 72°C were used. PCR products were purified with the MSB Spin PCRapace Kit (Stratec Molecular, Coronado CA, USA) and then Sanger-sequenced. Full-length consensus viral sequences were obtained assembling the sequences of the three amplified products by using the Genious R9.0.2 program.

### Construction of TuMV *VPg* mutants

Mutants in the *VPg* cistron were synthesized by long inverse site-directed PCR mutagenesis using the QuickChange II XL Kit (Stratagene, San Diego CA, USA) following the manufacturer’s instructions and using p35STunos as template. The mutagenic primers were designed following the manufacturer’s recommendations. Primer set cpr5-2F (5’-GGAGGATGAGTTGGGTCCAAATGAAATACGTGT-3’) and cpr5-2R (5’-ACACGTATTTCATTTGGACCCAACTCATCCTCC-3’) was used to generate the A6238G (D113G) mutant while primer set jin1F (5’-GGATCCAAATGAAATACATGTGAATAAGACAATTC-3’) and jin1R (5’-GAATTGTCTTATTCACATGTATTTCATTTGGATCC-3’) to obtain the G6253A (R118H) mutant. PCR protocol consisted of: a denaturation step of 2 min at 95 °C followed by 20 cycles of 20 s at 95 °C, 10 s at 60 °C and 7 min at 68 °C, and a final extension of 5 min at 68 °C. After *Dpn*I digestion and transformation of electrocompetent *Escherichia coli* DH5*α*, the presence of the desired mutations in the infectious clone and the absence of undesired nucleotide changes was confirmed by sequencing.

### Bioassays of the TuMV *VPg* mutants

The infectivity, viability and pathogenicity of both VPg TuMV mutants was confirmed by performing three independent bioassays. In each experiment, three batches of 24 *A. thaliana* plants (three-weeks old) of genotypes WT, *cpr5-2* and *jin1* were inoculated with the WT TuMV plasmid clone used as reference and with plasmids of each of the TuMV *VPg* mutants. Another two batches of 24 plants from *cpr5-2* or *jin1* genotypes were inoculated with the WT TuMV plasmid and the corresponding plasmid of the TuMV VPg mutant, the VPg/D113G mutant in the case of the *cpr5-2* genotype and that of VPg/R118H for the *jin1* genotype. *A. thaliana* plants were inoculated by abrasion of three leaves applying equal amounts of the plasmid inoculum, a total of approximately 7 μg per plant distributed in 3 μL per leaf. Plasmids were previously purified by Midiprep using the NucleoBond Xtra Midi Kit (Macherey-Nagel, Düren, Germany) and resuspended in distilled water. Number of infected plants and symptom intensity were collected every day until 15 dpi.

### Statistical analyses

All statistical analyses described hereafter were performed with SPSS version 27 software (IBM, Armonk, NY), unless otherwise indicated.

*AUDPS* (figure 2), *I* (figure 3) and *VL* (figure 4) data were fitted all together to a multivariate analysis of covariance (MANCOVA) model with plant genotype (*G*) as main factor, independent evolution lineages (*L*) were nested within *G* and passage (*t*) was introduced in the model as a covariable. The full model equation thus reads:

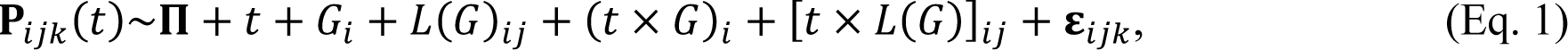

where **P***_ijk_*(*t*) = (*AUDPS_ijk_*, *I_ijk_*, *VL_ijk_*)^T^ is the vector of phenotypic traits observed at time *t*, for an individual infected plant *k* of evolutionary lineage *j* of genotype *i*, **Π** represents the vector of phenotypic grand mean values and **ε***_ijk_* stands for the vector of errors assumed to be Gaussian distributed at every *t*. The significance of each factor, the covariable and their interactions was evaluated using the Wilks’ *Λ* method. The magnitude of the effects was evaluated using the η^2^_P_ statistic (proportion of total variability in the traits vector attributable to each factor in the model; conventionally, values of η^2^_P_*≥* 0.15 are considered as large effects).

Rates of phenotypic evolution for *AUDPS* and *I* were estimated by fitting the time series data to a first-order autoregressive integrated moving-average, ARIMA(1,0,0), model (Elena and Sanjuán, 2005; González et al. 2019). The model equation fitted has the form:

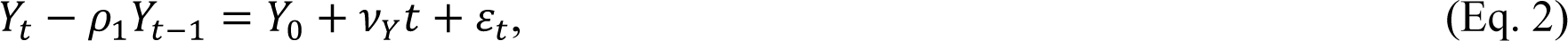

where *Y_k_* represents the variable being analyzed at passage *k*, *ρ*_1_ measures the degree of self-similarity in the time-series data (correlation between values at passages *t* and *t* – 1), *ε_t_* represents the sampling error at passage *t*, and *ν_Y_* represents the linear dependency of variable *Y* with passage number, that is, the rate of phenotypic evolution. The rates of phenotypic evolution were further analyzed in the context of the mutated defense signaling pathway genes or recessive resistances. Rates of phenotypic evolution were fitted to a multivariate analysis of variance (MANOVA) in which the plant genotype, serving as host for the evolutionary lineages (*G*), was nested within the factor being analyzed (*i.e*., the type of selection –hard *vs* soft-, the mode of resistance –signaling pathways *vs S*-, and whether mutations affected the SA-dependent, the JA/ET-dependent signaling or the RNA-silencing pathways; table 1). The full model equation now thus reads:

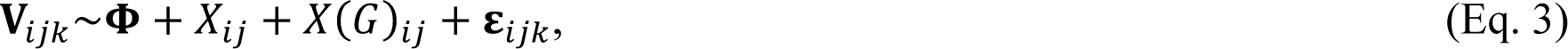

where **V***_ijk_* = (*ν_AUDPS_*, *ν_I_*)^T^ is the vector of rates of phenotypic evolution observed for the *k* replicate lineage evolved in plant genotype *j* within the factor *X_i_*, **Φ** represents the vector of grand mean values and **ε***_ijk_* the vector of Gaussian errors.

### Analysis of infection network

The first statistical approach consisted in fitting a logistic regression model to the presence/absence of symptoms data using GLM techniques with a Binomial probability distribution and a probit link function. The model equation reads as follows:

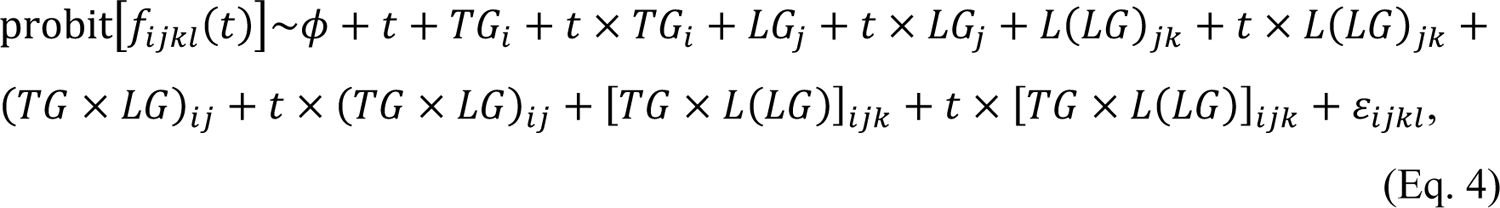

where *f_ijkl_*(*t*) is the frequency of symptomatic plants *t* dpi in the test host genotype *TG*, for the viral lineage *L* evolved in the local host genotype *LG*. *TG* and *LG* were considered as orthogonal factors, *L* was nested within *LG* and dpi (*t*) was treated as a covariable. The model includes all main factors, their corresponding nested and factorial interactions as well as their interactions with the covariable. *ϕ* represents the grand mean of the probit-transformed *f* values, and *ε* the error term assumed to be Binomial. The significance of the different factors was evaluated using likelihood-ratio tests (LRT).

For the second statistical approach, the binary infection matrix was analyzed using tools borrowed from the field of network biology to explore whether they show random associations between viral lineages and host genotypes, one-to-one associations, nestedness indicative of a GFG type of interaction, or modularity (Weitz et al. 2013). The statistical properties of the infection matrix were evaluated using the R package “bipartite” version 2.15 (Dormann and Strauss 2008) in R version 4.0.0 (R Core Team 2020) under RStudio 1.2.1335. Four different summary statistics were evaluated: *T* nestedness (Bascompte et al. 2003), *Q* modularity (Newman 2006) and the *d’* species-level (or Kullback-Leibler divergence) and $^#^network-level (or two-dimensional normalized Shannon entropy) specialization indexes (Blüthgen et al. 2006). $^#^ranges between zero and one for extreme generalists and specialists, respectively. Statistical significance of these statistics was evaluated using Bascompte et al. (2003) null model.

### The statistics of molecular evolution

Treating each lineage as an observation and each host genotype as a subpopulation, we evaluated the average nucleotide diversity within host genotypes, *π_S_*, the nucleotide diversity for the entire sample, *π_T_*, the interhost genotypes nucleotide diversity, *δ_ST_*, and the estimate of the proportion of interhost genotypes nucleotide diversity, known as coefficient of nucleotide differentiation (Nei 1982), *N_ST_* = *δ_ST_*/*π_T_*. Standard deviations of estimates were inferred from 1000 bootstrap samples. All these computations were done using MEGA 11 (Tamura et al. 2021) and the lowest-BIC nucleotide substitution model Kimura 2-parameters (Kimura 1980). Tajima’s *D* test of selection (Tajima 1989) and its statistical significance were evaluated using DnaSP6 (Rozas et al. 2017).

The frequency of mutations (*m*) per cistron (*C*), relative to the length of the corresponding cistron, was fitted to the following logistic regression model using GLM techniques with a Binomial probability distribution and a probit link function:

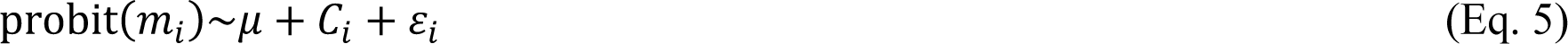

where *μ* is the average genomic mutation frequency and *i* refers to the 10 cistrons in the main ORF.

The functional effect of mutations found in the *VPg* cistron were evaluated *in silico* using the SNAP2 algorithm (Hecht et al. 2015). The algorithm provides a functionality score and an accuracy index for each possible mutation affecting a coding sequence. Negative score values mean functional neutrality while positive values should be taken as indications of functional effects; the larger the values, the stronger the effect.

## Supplementary Material

Supplementary data are available *at Molecular Biology and Evolution* online.

## Acknowledgments

We thank Francisca de la Iglesia and Paula Agudo for excellent technical assistance and the rest of the EvolSysVir lab members for fruitful discussions. This work was supported by grants BFU2015-65037-P and PID2019-103998GB-I00 (Agencia Estatal de Investigación - FEDER) and PROMETEU2019/012 (Generalitat Valenciana) to S.F.E.

